# Single-cell eQTL mapping reveals convergent glial–neuronal risk architecture in Parkinson’s disease

**DOI:** 10.64898/2026.04.24.720642

**Authors:** Zechuan Lin, Jacob Parker, Vanitha Nithianandam, Sakthikumar Mathivanan, Tao Wang, Zhixiang Liao, Sean K. Simmons, Idil Tuncali, Xian Adiconis, Nathan Haywood, Beatrice Weykopf, Xufei Teng, Monika Sharma, Jie Yuan, Clare Baecher-Allan, Xianjun Dong, Thomas G. Beach, Geidy E. Serrano, Joshua Z. Levin, Su-Chun Zhang, Mel B. Feany, Clemens R. Scherzer

## Abstract

Synucleinopathies affect ∼15 million people and are classically divided into neuronal (Parkinson’s disease (PD), dementia with Lewy bodies) and glial (multiple system atrophy) disorders. Here we challenge this dichotomy. We functionally fine-map 90 PD GWAS signals across nine cell types in cortex and substantia nigra using disease-context, population-scale single-nucleus eQTL meta-analysis (N = 1,197), bulk brain eQTL analysis (N = 1,182), and Mendelian randomization. A stringent causal framework integrates single-nucleus allelic imbalance (snASE) with orthogonal validation. We identify 125 functional risk genes for 50 loci—nearly doubling supported genes—and assign genes and cell types to over half of GWAS signals. Unexpectedly, 51% of risk genes are regulated in glia, particularly oligodendrocytes and their precursors. Across cell types, risk converges on a shared glial–neuronal vesiculopathy network. These findings uncover a convergent glial-neuronal risk architecture and establish a single-cell atlas for context-aware gene discovery and precision therapeutics for PD.

## Introduction

Neuronal alpha-synuclein diseases^1,2^—Parkinson’s disease (PD) and Lewy body disease—affect more than 15 million people worldwide^3,4^. Their pathological hallmark is the Lewy body, an intraneuronal aggregate found in dopaminergic neurons and cortical neurons^5-7^, composed of alpha-synuclein, vesicles, and dysfunctional mitochondria^8,9^. In contrast, the primary glial alpha-synuclein disease—multiple system atrophy—affects about a few hundred thousand people by some estimates^10^ and is characterized by glial cytoplasmic inclusions of alpha-synuclein^11^, rather than neuronal aggregates^12^.

PD is a synucleinopathy that classically affects the substantia nigra dopamine neurons during early disease stages causing impaired movements. However, it has become clear that PD is a multi-system, multi-region disease^7,13,14^, with widespread cortical involvement. Cortical Lewy body pathology is common across temporal, cingulate, and frontal cortices^7,13,14^ and is strongly associated with cognitive and psychiatric symptoms, including executive dysfunction, hallucinations, depression, and dementia^7,15-17^. Indeed, cortical Lewy body burden is the strongest neuropathological correlate of cognitive impairment in PD and a major determinant of quality of life and health care costs^7,13,18^. Emerging evidence further suggests that PD might affect multiple neuronal and glial cell types. Cortical glutamatergic neurons carry a substantial Lewy body burden that correlates with dementia^5-7^. Microglia^19,20^ and astrocytes^21-23^ are implicated in non–cell-autonomous alpha-synuclein–mediated neurotoxicity, and PD GWAS variants are enriched near oligodendrocyte-specific genes^24^, pointing to a broader cellular architecture of PD risk.

In the post-genome-wide association studies (GWAS)^25,26^ era systematic mapping of risk variants to their functional cellular effects is needed. GWAS loci often span large genomic intervals and encompass multiple genes, thus identifying the causal target genes and cell types, and resolving the functions of multiple signals within one locus requires approaches beyond simple proximity mapping.

Bulk-tissue eQTL^25-27^ and more recent single-nucleus eQTL studies have demonstrated the potential of functional fine-mapping in human cortex, particularly for Alzheimer’s disease and schizophrenia^28-35^, including work from our group^36^. Meta-analysis across these cohorts can partially mitigate their sample size and power limitations. For example, Ref.^37^ combined four of these cortex datasets (N = 757 individuals, largely Alzheimer’s disease and controls), modestly improving cell-type–specific eGene discovery for PD(e.g., 66 risk genes for 38 of >90 PD-associated GWAS signals).

However, none of these studies included PD brain samples, which are likely essential given that GWAS variants are thought to act through cell-type–specific regulatory mechanisms shaped by disease state and cellular vulnerability^7-32^. As a result, prior efforts have had limited ability to assign functional risk genes to the majority of PD GWAS loci.

A comprehensive, cell-type–resolved atlas that functionally maps Parkinson’s disease (PD) GWAS variants to causal genes, target cell types, and biological pathways across vulnerable dopaminergic and cortical circuits has remained elusive. Prior studies have been limited by underrepresentation of disease-relevant cell states, insufficient power to resolve cell-type–specific regulatory effects and complex loci, and a lack of rigorous causal validation, particularly in the most vulnerable neuronal populations. Such limitations have hindered a systematic understanding of PD genetic architecture and the identification of actionable therapeutic targets.

Here we present PD5D+, a high-resolution functional atlas of PD genetics that overcomes these constraints by integrating disease-contextualized, cell-type–specific regulatory maps with large-scale meta-analytic power and orthogonal validation. We performed single-nucleus eQTL analyses in 94 human brains spanning prodromal, motor, and cognitive stages of PD, explicitly capturing disease-relevant regulatory states. To maximize statistical power and generalizability, we integrated these data with massively scaled meta-analyses of single-cell (n = 1,197) and bulk substantia nigra and cortical eQTL datasets (n = 1,182), totaling 2,279 unique human brains. Critically, our design includes dopaminergic neurons—the most vulnerable cell type in PD—and cortical cells from PD patients, enabling, for the first time, cell-type–specific eQTL mapping directly within the PD disease context.

We further establish a rigorous causal inference framework by validating *cis*-regulatory effects at single-nucleus resolution using allelic imbalance analyses (snASE) in heterozygous individuals, complemented by comprehensive orthogonal molecular and functional assays. This integrative strategy nearly doubles the number of functionally supported PD risk genes to 125 and assigns target genes and cell types to more than half of PD GWAS loci. Strikingly, PD risk converges on a shared glial–neuronal vesiculopathy network, challenging the traditional neuron-centric view and revealing a continuous risk architecture across cell lineages. Mechanistically, we identify *BIN3* as the functional effector of the Chr. 8q21.3 PD GWAS locus and demonstrate that its dysregulation perturbs vesicle cycling and exacerbates α-synuclein toxicity, directly linking genetic risk to disease biology.

## Results

### Discovering cell type eQTLs associated with noncoding GWAS variants of Parkinson’s disease in 1,197 individuals and 5.6 million single human brain cells

#### Functional fine-mapping ninety genetic loci associated with Parkinson’s to their cellular functions (Fig. 1)

Gene expression is the most proximal function of the genome. Here, we mapped the functions of **32,267** PD-associated genetic variants representing 90 GWAS peaks (78 loci) at scale and cellular resolution in nine disease-relevant cell types --- dopamine neurons, glutamate neurons, GABA neurons, oligodendrocytes, oligodendrocyte precursor cells (OPCs), astrocytes, microglia, endothelial cells, and pericytes --- from substantia nigra and cortex of a total of **1,197** individuals. We analyzed our PD5D (this study) and BRAINcode cohorts^38-40^ (**Supplementary Table S1 and Supplementary Table S2**) and enhanced statistical power through meta-analysis^31,32,41^. PD5D and BRAINcode^38-40^ span healthy individuals and Parkinson’s patients across the entire disease-span. They include onset disease --- hard-to-find neurologically normal individuals, whose brain autopsy revealed hallmark prodromal, PD-associated Lewy neuropathology with alpha-synuclein positive Lewy body Braak stages I-III^14^ --- as well as PD patients with more advanced cortical Lewy body stages (**Supplementary Table S1 and Supplementary Table S2**). Statistical power for detecting the nine cell type-eQTLs was increased through large-scale meta-analysis including single-cell data sets of human cortex^31,32^ and of mesencephalic dopamine neurons derived from human controls’ iPSCs^41^.

**Fig. 1.**
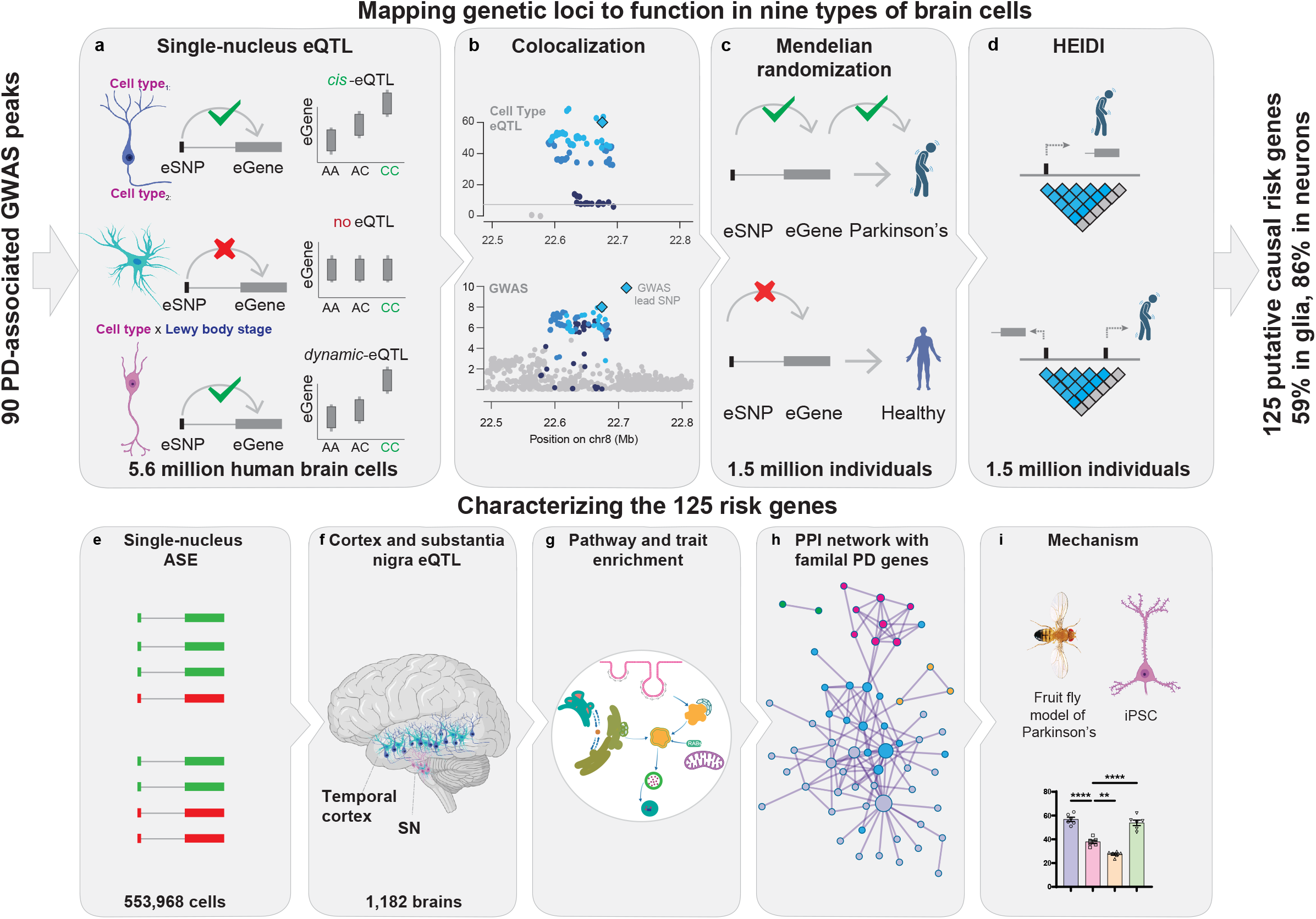
Fine-mapping ninety PD loci to their cellular functions: study design. We analyzed gene regulation across nine cell types from substantia nigra and cortex of 1,197 individuals—two brain regions central to the motor and cognitive manifestations of PD. Putative causal PD risk genes were prioritized using four criteria: **a**, Cell-type–specific eQTL analysis in 1,197 subjects, encompassing ∼5.6 million human brain cells from cortical and midbrain tissues. **b**, Nomination of PD risk genes by colocalization of cell-type–specific cis-regulatory eSNPs with PD GWAS variants (posterior probability ≥80%). **c**, Confirmation of PD risk genes by causal associations between gene expression and PD risk using SMR. **d**, Confirmation of PD risk genes using the HEIDI test to distinguish shared causal variants from linkage between GWAS and eQTL signals. In total, 125 genes met all criteria. These genes were further characterized by multiple molecular assays: **e**, Large-scale validation of PD risk genes by single-nucleus allele-specific expression (sn-ASE) in 553,968 brain cells. **f**, Recovery of eQTL associations in cortex and substantia nigra homogenates from an independent cohort of 1,182 individuals. **g**, Pathway and disease-trait enrichment analyses providing orthogonal evidence of PD association. **h**, Protein–protein interaction (PPI) network analysis incorporating familial PD genes, providing additional orthogonal support. **i**, Functional testing of a prioritized PD risk gene in a *Drosophila* PD model and in human iPSC-derived neurons. Overall, 102 of the 125 putative causal risk genes were supported by at least one additional line of evidence.

**Fig. 2.**
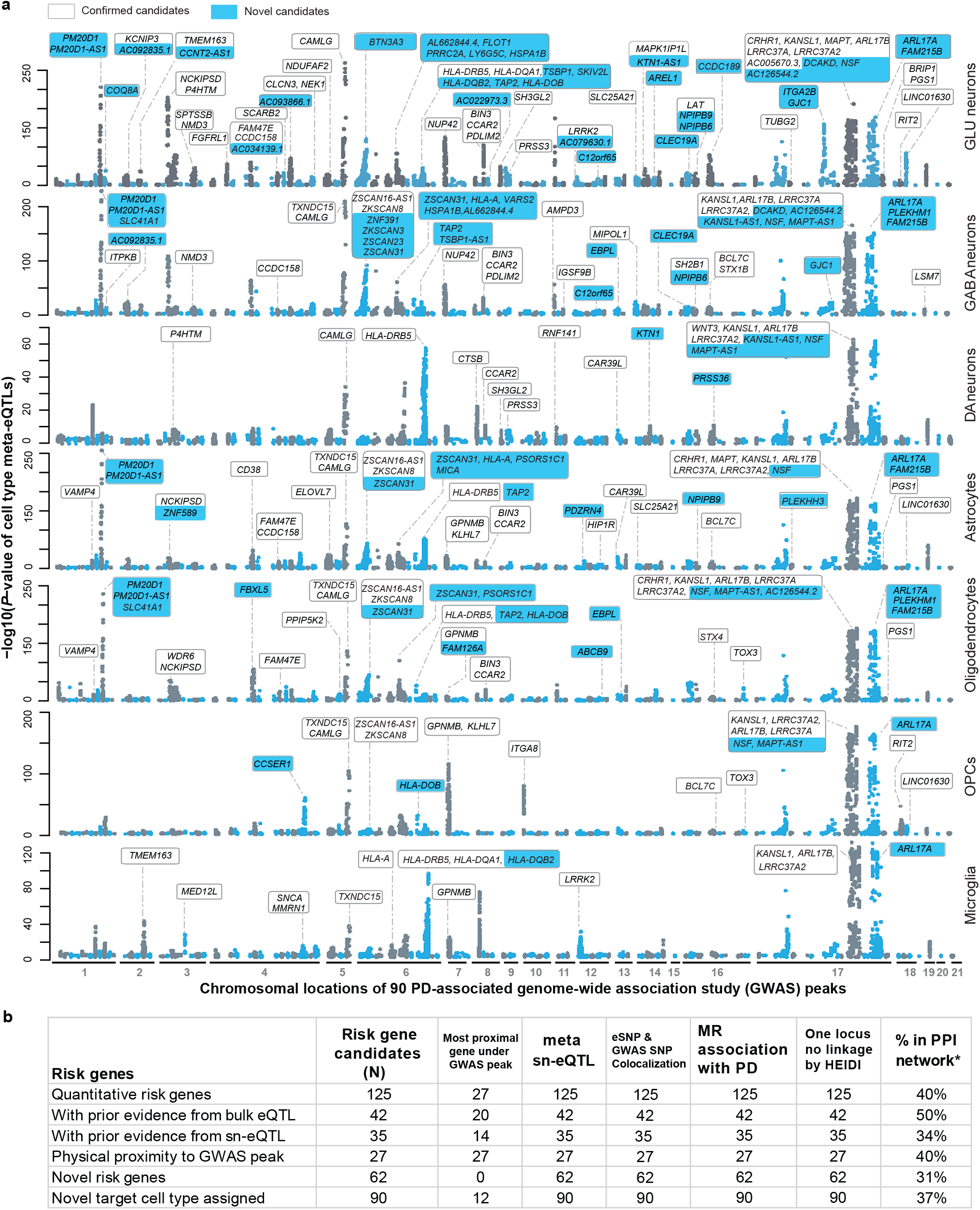
125 quantitative genes associated with PD risk in specific brain cells. **a**, We fine-mapped ninety PD loci to cellular functions across nine brain cell types and 5.6 million single cells using cell type–eQTL meta-analysis in 1,197 individuals. Manhattan plots show associations between variants in ninety GWAS peaks and gene expression in each cell type (Y-axis: –log_10_ P; X-axis: chromosomal positions). We identified 125 cis-regulated putative causal risk genes, including novel genes (cyan), and mapped their target cell types. Previously nominated risk genes appear on a white background. **b**, The table summarizes fine-mapping evidence for all 125 genes (columns) and compares them to prior bulk eQTL, sn-eQTL, and physical mapping efforts. The study identifies 63 novel risk genes and assigns 98 genes to new target cell types.

We conservatively required **four rigorous significance requirements** to prioritize putative causal PD-associated risk genes (**Fig. 1a-d**): i, significant variant-gene expression association in a cell type indicating that DNA variants (eSNPs) regulate that gene’s (eGene) transcription (cell type-eQTL). ii. colocalization of the implicated *cis*-regulatory eSNPs and GWAS risk variants with 80% or higher posterior probability; iii. evidence for a causal association between eGene expression and PD risk using Summary-based Mendelian randomization; iv. no evidence of linkage by Heterogeneity in Dependent Instruments (HEIDI) test consistent with a true shared causal variant. The HEIDI test distinguishes true shared causal variants from linkage by testing whether a GWAS signal and an eQTL signal arise from the same underlying variant rather than from two correlated but separate variants^42^.

Temporal cortex and substantia nigra brain samples were obtained through a rapid autopsy program with superb post-mortem intervals (median PMI, 3 h) and RNA quality (median RIN, 9.2) for **94** and **83** individuals (**Supplementary Table S1**), respectively. *Cis*-eQTL analyses were performed for 32,267 PD-associated variants—including 7,057 GWAS variants from 90 loci, 3,550 sub-threshold GWAS variants (P ≤ 10^−6^), and 21,660 linked proxy variants (with r^2^ ≥ 0.4)— in glutamate neurons, GABA neurons, oligodendrocytes, OPCs, astrocytes, microglia, endothelial cells, pericytes using single-nucleus RNA-seq and computational pseudobulking (see Methods). Dopamine neurons, comprising only 0.5–2% of midbrain cells, pose challenges for single-nucleus RNA-seq. Enrichment strategies based on FACS-sorting from substantia nigra are constrained by limitations of nuclear markers (e.g. NeuN is not reliable for dopamine neurons^43^; Nurr1 appears biased to dopamine neurons without alpha-synuclein pathology and without aging-changes^44^). To overcome cell rarity and selection biases inherent to enrichment methods, we performed laser-capture RNA-seq (lcRNA-seq)^40^ of neuromelanin-positive dopamine neurons from human substantia nigra, experimentally bulking 300–800 neurons per substantia nigra pars compacta of **83** controls and individuals with onset and advanced PD-associated Lewy body neuropathology from incipient brainstem to advanced cortical Lewy body Braak stages as detailed in Ref.^40^ (**Supplementary Table S2**). The power and validity of lcRNA-seq for detecting even low abundance regulatory RNAs (e.g. long noncoding RNAs, enhancer, and circular RNAs) with high cell-type specificity and without compromising anatomical or morphological context was previously demonstrated^38-40^. Models were adjusted for biological and technical covariates (Methods). To maximize statistical power, analyses focused on *cis*-regulatory effects.

#### Cell type-eQTL meta-analysis (Fig. 1, Fig. 3a)

To enhance statistical power for detecting cell type-eQTLs for low-frequency cell types and small effect sizes, we increased sample sizes through meta-analysis of a total of 1,197 unique individuals and 5.6 million cells across the current PD5D data set, BRAINcode^38-40^, sn-eQTL datasets from cortex of 905 individuals^31,32^, and sn-eQTL data from mesencephalic dopaminergic neurons of 215 individuals^41^ (**Supplementary Table S2**). We discovered **271** *cis*-regulated genes in nine cell types with *P* values equal or below (more significant) the conservative Bonferroni threshold of 3.38 × 10^-7^ indicating significance (e.g. 0.05/the number of 147,929 unique eSNP-eGene pairs; **Supplementary Table S3)**. To reveal independent, secondary *cis*-regulatory associations, we performed conditional LD clumping and LD-chunk analysis (Methods), elucidating 324 additional eQTLs. For 127 of these eGenes multiple independent eQTLs were detected corresponding to a total of 595 cell type-eQTLs (595 unique eSNP-eGene pairs representing 271 eGenes and 514 eSNPs). The 271 eGenes were cis-regulated by 73 of the 90 PD-associated GWAS peaks.

**Fig. 3.**
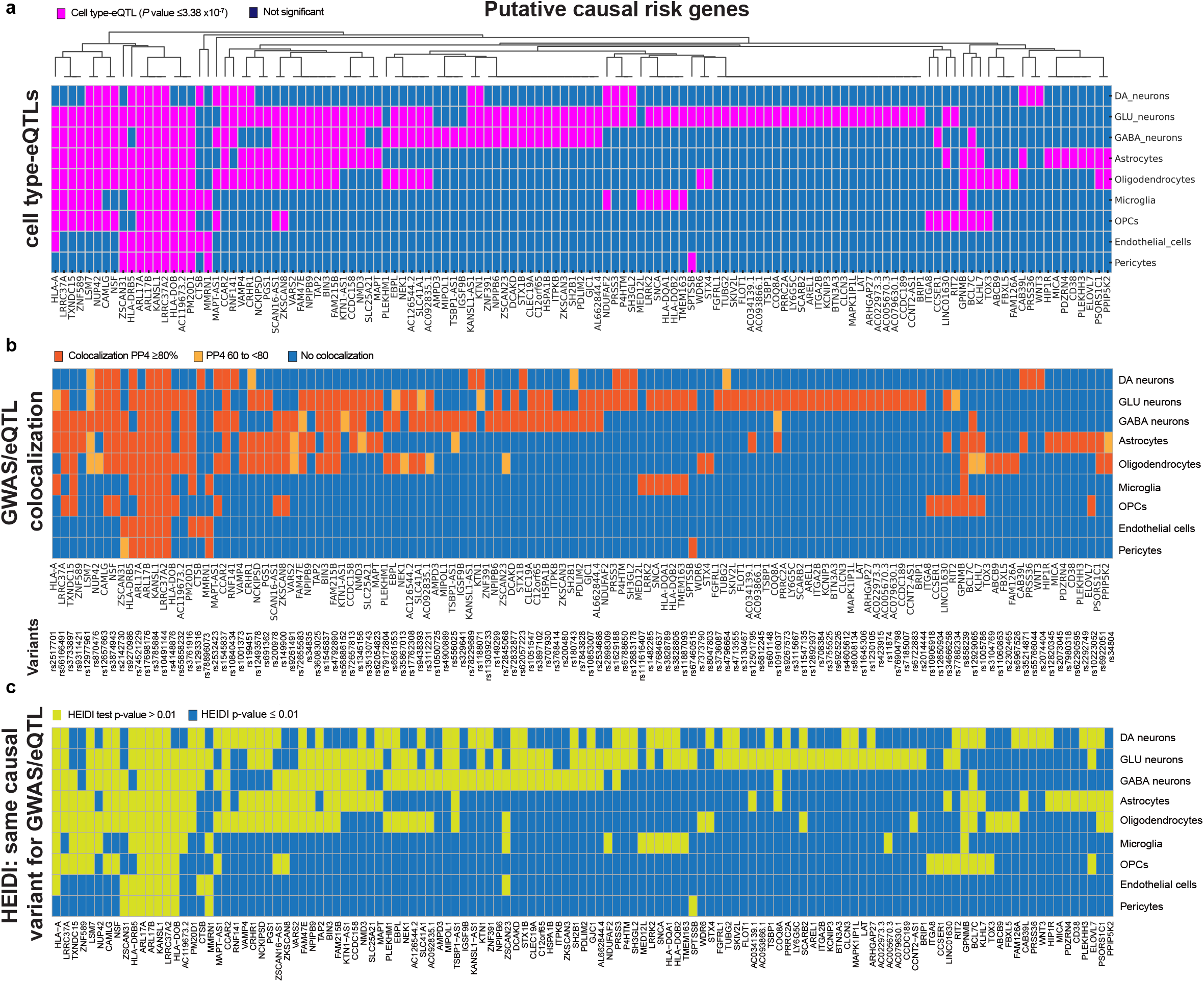
Prioritizing putative causal PD risk genes using cell type–eQTL meta-analysis, colocalization, causality, and linkage tests. **a-c**, Risk genes were required to meet four stringent criteria: significant cell type–eQTLs showing that variants (eSNPs) regulate gene expression (eGenes; **a**, magenta cells, eQTLs passing the 3.38 × 10^-7^ threshold); colocalization between cis-regulatory eSNPs and GWAS variants with ≥80% posterior probability (**b**; orange cells); causal support from summary-based Mendelian randomization (Supplement); and no evidence of linkage in the HEIDI test (**c**; lime cells, HEIDI *P* > 0.01), consistent with a single shared causal variant. Matrices show the 125 prioritized genes (columns) across nine cell types (rows).

#### Colocalizing expression and trait variants (Fig. 3b)

We employed the coloc test^45^ which uses Bayesian statistic to test the *a priori* assumption of whether a single causal variant may be regulating both target gene expression and disease risk. We evaluated the 595 unique SNP-gene pairs (representing 271 eGenes and 514 eSNPs) forwarded from the cell type-eQTL analysis. For 156 eGenes, colocalization analysis of eSNPs and GWAS SNPs indicated high (≥0.8) posterior probability (PP4) for a single shared causal variant explaining the association with both the expression and disease trait (**Supplementary Table S4**). PP4 equal or greater than 0.8 was conservatively used as evidence for a single variant associating with both gene expression and PD risk^45^.

#### Causality test

To identify which cell type–specific quantitative trait genes may be causally associated with Parkinson’s disease, we applied Mendelian randomization. Using cis-regulatory eSNPs associated with expression of the 271 eGenes as instrumental variables, we tested for associations with PD in summary statistics from 37,688 PD cases and 18,618 proxy cases compared to 1.4 million controls^46^. Proxy cases are individuals without PD who have a first-degree relative with PD and are used to capture inherited genetic risk^46^. Nalls et al.^46^ and we performed extensive sensitivity analyses and found no material influence of proxy cases on the lead SNPs used for downstream functional analyses (see Discussion). A total of 211 eGenes achieved *P*-values at or below the Bonferroni threshold for significance (1.8 × 10^−4^; 0.05/271 eGenes; **Supplementary Table S5**), indicating associations between eGene expression and PD under an assumption of causality or pleiotropy^42^.

#### Heterogeneity test (Fig. 3c)

We next assessed whether the same causal variant influences both gene expression and PD risk using the Heterogeneity In Dependent Instruments (HEIDI) test^42^ that distinguishes a shared causal variant from linkage between eQTL and GWAS signals. A non-significant HEIDI test supports a shared causal variant, whereas a significant HEIDI test suggests heterogeneity consistent with linkage between distinct variants. All prioritized candidates showed non-significant HEIDI P-values (>0.01), consistent with a single causal variant underlying both the gene-expression and PD-trait associations.

In summary, 125 putative causal risk genes and 124 putative causal risk variants across 50 PD-associated GWAS peaks met our stringent four-tier significance criteria in at least one of the nine cell types. These criteria included: (i) sn-eQTL P values below 3.38 × 10^− 7^; (ii) colocalization of eSNPs and GWAS SNPs; (iii) association with PD risk in Mendelian randomization; and (iv) a non-significant HEIDI test supporting a shared causal variant. Based on this convergent evidence, we prioritize these 125 genes as putative causal quantitative risk genes associated with common, “idiopathic” PD (**Supplementary Table S6**).

### Characterizing prioritized risk genes with nine molecular modalities

**We further characterized these candidates through nine molecular analyses** (**Fig. 1**), including i. **sn-ASE analysis**, ii. **cortex- and substantia nigra**-**eQTL analysis**, iii. **dynamic sn-eQTLs of Lewy body burden**, iv. **pathway** and v. **disease trait** enrichment, vi. **protein-protein network** interactions with familial PD genes, vii. **loss-of-function analyses**, and viii. performed **proof-of-concept mechanistic testing in human PSC-derived neurons** and ix. a ***Drosophila* model of alpha-synucleinopathy**.

#### PD-associated variants function, in part, through allele-regulatory effects tailored to cell types (Fig. 4)

We evaluated *cis*-regulatory effects at the level of allele-specific expression *in vivo* in human brain cells by deploying for the first time at scale single-nucleus allele-specific expression (sn-ASE) analysis, an experimental-computational method we recently developed^47^, for 550,450 human single brain nuclei **(Fig. 4)**. sn-ASE pinpoints the precise effect of each disease-linked vs. reference allele on phased reads and *allele-specific transcripts within* each *individual brain cell* from *heterozygous* individuals, while eQTL addresses genetic effects on *gene expression in-between groups of individuals with different genotypes*. By evaluating ASE within each cell, power is increased as technical and biological artifacts are controlled for^48^. We used phased genotypes and a pseudobulk ASE dataframe (pseudobulked by sample of origin and cell type; see Methods for detail). We tested for ASE difference between alleles by fitting a beta-binomial generalized linear model on this pseudobulked ASE data for each cell type. Only variants with ≥ 20 phased UMIs were retained in the study. 77 out of the 125 GWAS target eGenes had sufficient heterozygous individuals and phased reads available for this analysis (see Methods). Effect sizes from sn-ASE and sn-eQTL analyses were significantly concordant with *P* value = 1.08 × 10^-15^ and Pearson correlation coefficient of 0.61 (**Fig. 3**). For 48 of the 77 putative causal risk genes (62%) we detected ASE with *P* values ≤ 0.05 (**Supplementary Table S7**). For 52% of the target cell type-target gene pairs implicated by sn-eQTL analysis, *cis*-regulation of risk allele expression was observed as a plausible underlying molecular mechanism. The true number is expected to be even higher because sn-ASE analysis was limited to a subset (83 of 94) of individuals.

**Fig. 4.**
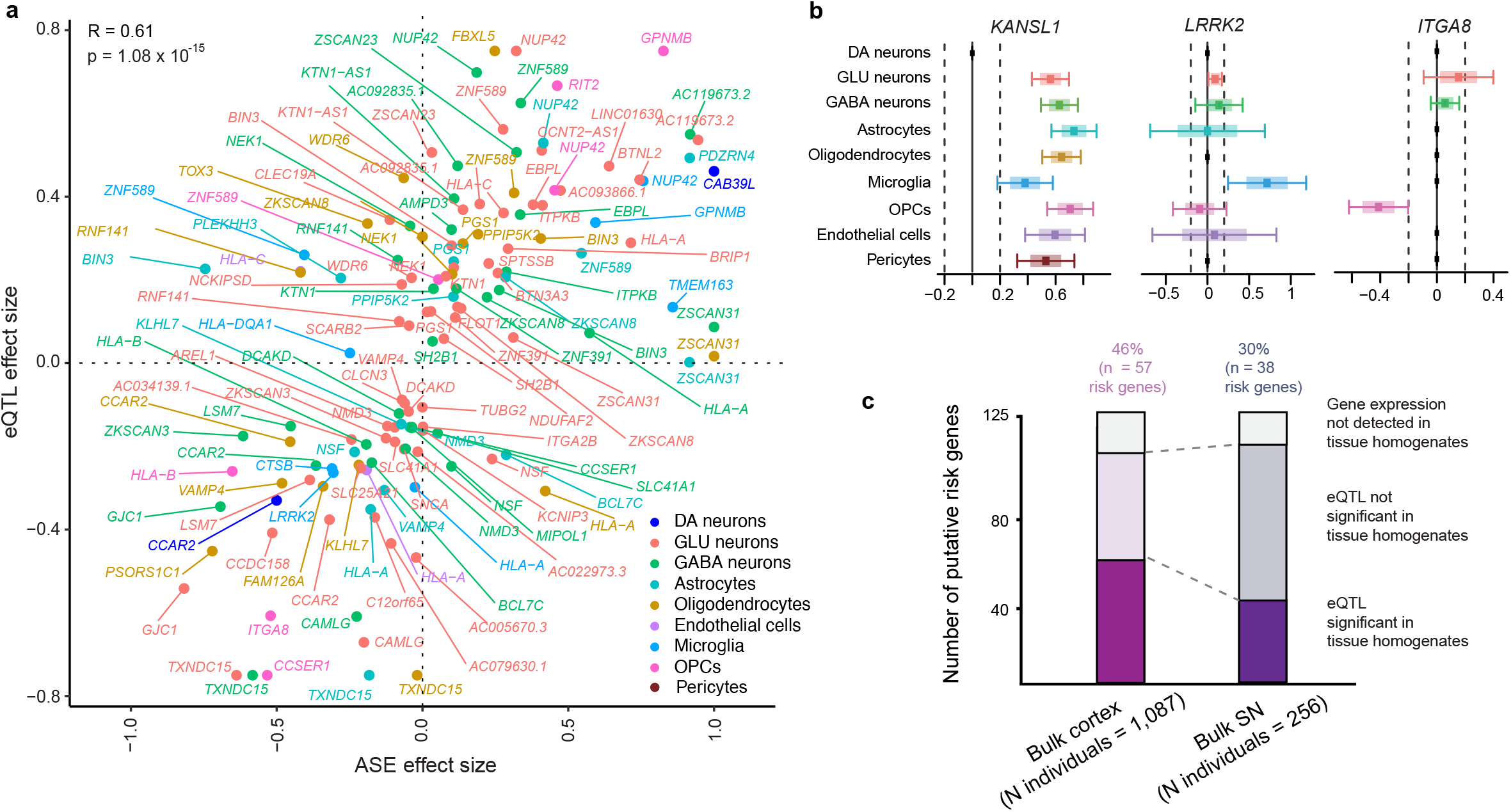
Single-nucleus allele-specific expression (sn-ASE) confirms sn-eQTLs. **a**, eQTL effects were validated by comparing counts of variant versus wild-type alleles in single brain cells using sn-ASE. sn-eQTL and sn-ASE effect sizes were significantly correlated (*P* = 1.08 × 10^−15^, R = 0.61). **b**, Box plots show allele-specific expression for *KANSL1, LRRK2*, and *ITGA8* in specific cell types. sn-ASE confirmed *cis*-regulation of *KANSL1* in 8 of 9 cell types (*P* < 5.3 × 10^−7^), *LRRK2* in microglia (*P* = 8.5 × 10^−5^) and glutamate neurons (*P* = 0.01), and *ITGA8* exclusively in OPCs (*P* = 2.7 × 10^−7^), consistent with sn-eQTLs. The x-axis shows the proportion of phased UMIs mapping to the GWAS effect allele.

#### Comparison to PD-associated gene-regulation in cortex and substantia nigra homogenates of another 1,182 independent individuals (Fig. 4c)

Single-nucleus eQTL increases the signal-to-noise ratio for identifying *cis*-regulatory mechanisms that are fine-tuned to the biology of a cell type, particularly if that cell type is under-represented in the tissue homogenates. Bulk RNAseq analyses assay expression changes in an admixture of multiple cell types without cell type resolution. Bulk RNAseq data sets however have the advantage of very large accessible sample sizes and resulting statistical power improvements. Thus, it is reasonable to expect that these large sample sizes can in part overcome the lack of cell type resolution, and that cell type-eQTLs affecting all or most cell types to be largely recoverable and those manifest in specific cell types to be partially recoverable from large-scale bulk eQTL analyses of the appropriate source regions from which the pertinent cell types were captured. We evaluated the 125 risk genes for PD identified in cell type-eQTL analyses in a large, population-scale eQTL analysis of human brain samples reported today from **1,182 unique individuals** representing seven cortex cohorts and two substantia nigra cohorts (**Supplementary Table S8**). In these populations the source regions for the nine cell types --- cortex and substantia nigra --- were homogenized and analyzed by bulk RNA sequencing or bulk gene expression microarrays. We evaluated brain cortex from 1,087 individuals from seven cohorts and in substantia nigra from 256 individuals from two cohorts (representing in total 1,182 unique individuals; **Supplementary Table S8**). 114 out of the 125 risk eGenes and 113 risk eSNPs were available for analysis in these region-specific populations. Eleven risk genes and eleven risk eSNPs were not available for this analysis due to differences in gene expression and genotyping platforms. Overall, *cis*-regulatory associations were recovered for 58 out of the available 114 risk genes (51%) with false discovery rates (FDR) ≤ 0.05. This is consistent with prior studies comparing cell type-eQTLs to bulk tissue-eQTLs (29) (**Supplementary Table S3**). As expected, recovery of cell type-eQTLs affecting many cell types (e.g. 8-9 cell types) was very high at 100% (4 of 4; e.g. the Chr. 17 risk genes, *KANSL1, ARL17A, ARL17B;* and Chr. 1 risk gene *PM20D1*). Moreover, eQTLs exclusively found in high-abundance cell types, e.g. oligodendrocytes, replicated at higher rates (43%) than eQTLs exclusively found in low-to-medium abundance cell types (e.g. 11% and 29% of eQTLs exclusively found in microglia and OPC, respectively).

#### Exploring the influence of local Lewy body burden on variant-gene expression associations (dynamic sn-eQTLs)

It is possible that in addition to cellular context, disease-context influences sub-sets of genetic gene-regulatory effects in PD. To begin to explore, whether subsets of the quantitative GWAS variant–transcript traits here identified may be activated or deactivated in response to accumulating alpha-synuclein-positive Lewy neuropathology, we evaluated the interaction of Lewy body score and GWAS variants on gene expression. We found suggestive evidence for possible interactions between Lewy body score and PD risk variants in regulating the cellular expression of 13 of the 125 quantitative trait genes nominal *P* values ≤ 0.01 (**Supplementary Table S9**). These initial clues warrant further evaluation in larger autopsy populations that were carefully neuropathologically phenotyped with semiquantitative Lewy body scores.

In parallel, **loss-of-function (LoF) burden tests** from a systematic rare-variant analysis of 7,184 PD cases, 6,701 proxy cases, and 51,650 controls^49^ and Ref.^25^ revealed LoF mutations for 18 of the 113 risk genes available for this analysis with nominal *P* values ≤ 0.05 (**Supplementary Table S10**). These two LoF investigations recovered ten genes supported both by our study and external datasets (*LRRK2, SCARB2, SH3GL2, CCAR2, CLCN3, CRHR1, KCNIP3, LAT, MMRN1, PPIP5K2*); and eight genes uniquely implicated by our analysis (*FAM126A, CCDC158, HLA-DOB, PDZRN4, SLC25A21, TSBP1-AS1*, and *ZSCAN31*). The number of true LoF-confirmed genes is likely higher, given that LoF variants are typically rare and our eQTL analyses excluded variants with minor allele frequency ≤ 5% to control statistical inflation.

#### The quantitative trait gene *BIN3*, regulated by the Chr. 8q21 locus (Fig. 7a-g), mechanistically enhanced activity-dependent endocytosis and alpha-synuclein aggregation (Fig. 7h-o)

BIN3 encodes a BAR-domain protein involved in sensing membrane curvature and interacting with small GTPases^50^. PD risk variants were associated with increased *BIN3* expression. Inducible *BIN3* overexpression significantly increased activity-dependent endocytosis in human stem-cell–derived neurons (**Fig. 7h-o and Fig. S2**). *BIN3* overexpression enhanced uptake of transferrin (Alexa Fluor 647)^51^ and increased FM1-43 FX fluorescence^52^. Increase in expression of the *BIN3* ortholog Amph in a *Drosophila* model of alpha-synucleinopathy^53^ worsened alpha-synuclein–induced locomotor deficits (**Fig. 7h**), enhanced neuronal loss (**Fig. 7i,j**), and alpha-synuclein inclusions (**Fig. 7k,l**) without intrinsic toxicity in the absence of human alpha-synuclein.

#### In summary, multiple lines of supporting evidence further substantiated 102 of the 125 prioritized risk genes

**(Fig. 1; Supplementary Table S10)**, including support from sn-ASE (48 genes; **Fig. 4**), independent bulk eQTLs across seven cortical and two substantia nigra cohorts (58; **Fig. 4**), dynamic-eQTL influenced by Lewy body score (13; **Supplementary Table S9**), disease gene enrichment analysis (**Fig. 5**; 7 neuronal risk genes (*SCARB2, MAPT, WNT3, TMEM163, STX1B, LRRK2*) annotated to sporadic PD; 4 glial risk genes annotated to MSA (*MAPT, SNCA, ELOVL7, LRRK2;* **Fig. S1**), protein–protein interactions with familial PD genes (45; **Fig. 5**), loss-of-function burden tests from Refs.^25,49^ (18), and mechanistic studies for one prioritized candidate (**Fig. 7**). In total, 102 of the 125 putative risk genes are supported by at least one additional line of evidence.

**Fig. 5.**
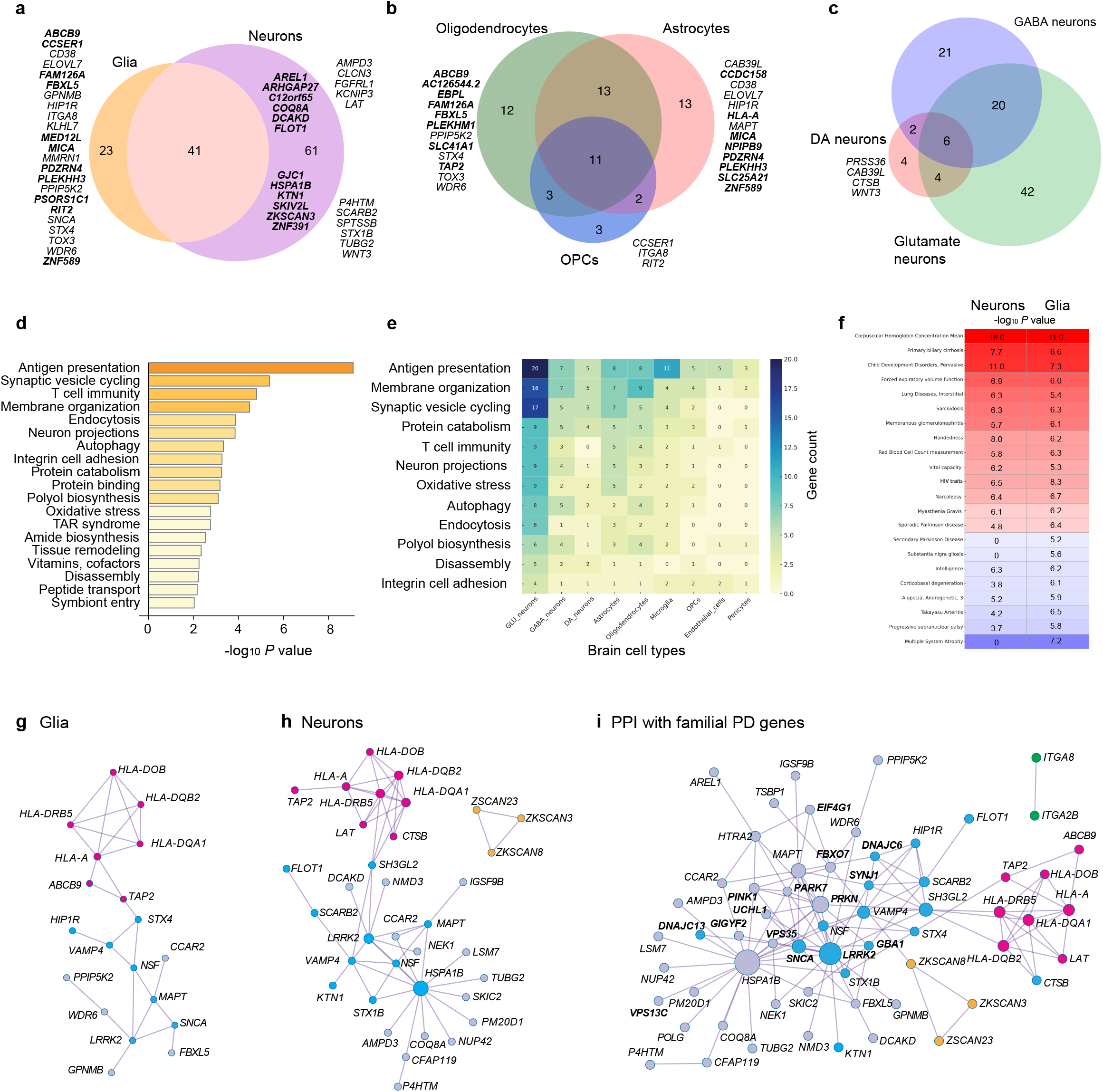
Risk genes converge on connected vesicle-cycling and antigen presentation networks contextualized to glia and neurons. **a–c**, Risk gene intersections in glia and neurons. **a**, Venn diagram of glial vs. neuronal risk genes; **b**, overlap among oligodendrocyte, astrocyte, and OPC risk genes; **c**, overlap among dopamine, glutamate, and GABA neuronal risk genes. Novel genes are in bold. **d–e**, Biological pathways and disease traits enriched in the 125 risk genes. **d**, Pathway enrichment (x-axis: –log_10_P, Fisher’s exact test). **e**, Heatmap showing the number of dysregulated risk genes per pathway (rows) across cell types (columns). **f**, Disease traits enriched in neuronal (left) and glial (right) risk genes. **g–h**, Protein–protein interaction networks for glial (**g**) and neuronal (**h**) risk genes. **i**, Combined PPI network of all 125 risk genes with 21 familial PD genes (bold font), revealing direct interactions with *DNAJC6, SYNJ1, PRKN, GBA1, LRRK2, SNCA, VPS35, VPS13C, PARK7, FBXO7, PINK1, GIGFY2,EIF4G1, UCHL1*. Blue circles mark vesicle-cycling genes; magenta marks antigen-presentation/loading genes. *HSPA1B* and *LRRK2* are major hubs connected to vesicle cycling.

#### Glial and neuronal cells mediating PD risk

Novel target cell types were assigned to 98 risk genes without prior sn-eQTL information. 51% of risk genes (64 of 125) were *cis*-regulated in glia (**Fig. 5a**), 80% in neurons (101 of 125), and 7% (9 of 125) in vascular cells (pericytes, endothelial cells). 69% of glial risk genes were significantly regulated in oligodendrocytic lineage cells (e.g. oligodendrocytes or OPCs), 63% in astrocytes (**Fig. 5b**), and 23% in microglia. Neuronal risk genes highly overlapped between glutamate, GABA, and dopamine neurons (**Fig. 5c**). The number of neuronal genes achieving all four significance criteria in dopamine neurons is likely an underestimation due to the limited sample sizes available for this rare cell type and power limitations.

#### Risk genes identification

62 of the risk genes resolved here at cell-type–specific regulatory resolution were not previously identified in earlier sn-eQTL studies^28-35^. Most were not the nearest gene to the lead GWAS variant (78%). Nearly half of the identified genes represent previously unrecognized, novel PD risk genes (**Fig. 2b, Table 1, Supplementary Tables S6 and S10**).

**Table 1.**
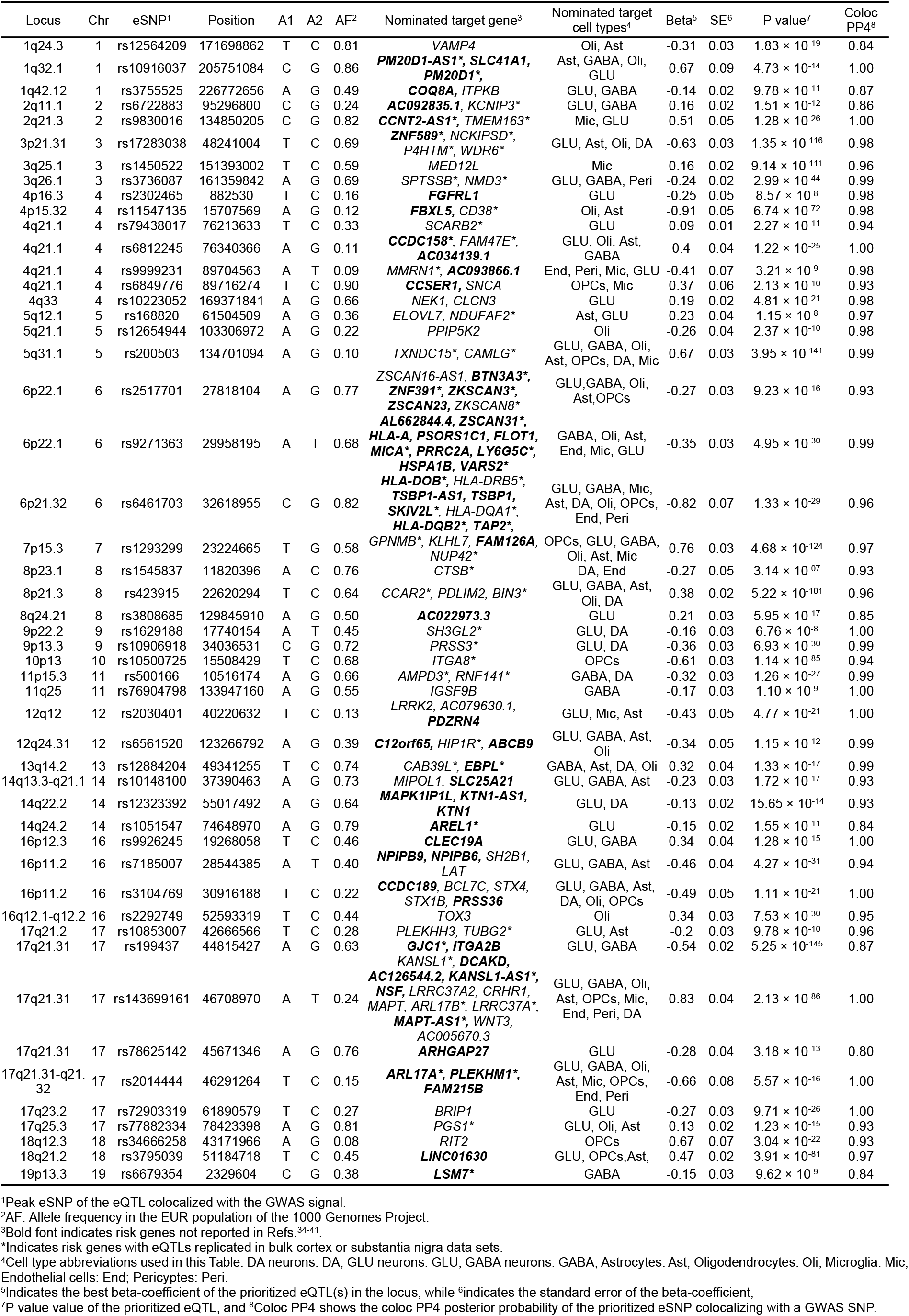
Identification of 125 PD-associated risk genes with cell type resolution in fifty GWAS peaks.

#### Cell context–aware regulation of antigen presentation machinery in glia and neurons (Fig. 5d-i)

Twenty-seven quantitative PD risk genes (22%) mapped to antigen presentation gene sets (**Fig. 5d,e**), forming a 17-gene PPI network (**Fig. 5i**). In the CNS, HLA class I genes show low basal expression in neurons and glia but are strongly induced by inflammation, enabling antigen presentation, while HLA class II genes are thought to be restricted largely to microglia^54^. Although oligodendrocytes are not classical antigen-presenting cells, multiple sclerosis and viral infections can activate MHC expression^55-57^. Peptide loading onto MHC molecules is mediated by the MHC loading machinery, including *TAP2* and *ABCB9*^58^, with *ABCB9* exclusively regulated in oligodendrocytes (**Fig. 5b,g**). *TAP2* linked antigen loading to vesicle cycling via *STX4* in glia (**Fig. 5g**). Loaded MHC molecules then trafficked through endosomal pathways to the plasma membrane. Regulatory effects on *TAP2* and *HLA-DOB* may be modulated by Lewy body burden (dynamic sn-eQTLs; **Supplementary Table S9**).

#### Twenty-five quantitative PD risk genes in glial and neuronal vesicle cycling

Approximately 20% of PD risk genes mapped to synaptic vesicle recycling and endocytosis pathways and formed a PPI network with familial PD genes (*DNAJC6, SYNJ1, GBA1, SNCA, LRRK2;* **Fig. 5i**).

#### Vesicle-cycle perturbations in glia remain largely unexplored (Fig. 5d,e)

PD GWAS variants quantitatively regulated 17 glial vesicle-cycle genes involved in SNARE assembly and membrane trafficking (*SNCA, VAMP4, BIN3* (**Fig. 7**), *LRRK2* (**Fig. S1**), NSF, *RIT2, STX4, HIP1R, NCKIPSD,CCDC158, CCAR2, WDR6, MAPT, CD38, SLC41A1, FAM126A*). In glia, vesicle cycling supports membrane remodeling and secretory programs distinct from neuronal neurotransmission: oligodendrocytes use VAMP7-positive late endosomal pathways and PI4KIIIα–FAM126A complexes to deliver myelin^59,60^; astrocytes release gliotransmitters^61^; and microglia deploy vesicles for cytokine secretion and antigen presentation^62^. *FAM126A* emerged as a novel PD gene in oligodendrocytes, consistent with its role in generating PI4P-rich vesicle-docking domains^60^. *EBPL*, a putative causal gene at the Chr13q14 peak, was regulated in oligodendrocytes and neurons and may influence cholesterol-dependent myelin biology based on homology to *EBP*^63^ (**Fig. 6a-f**).

**Fig. 6.**
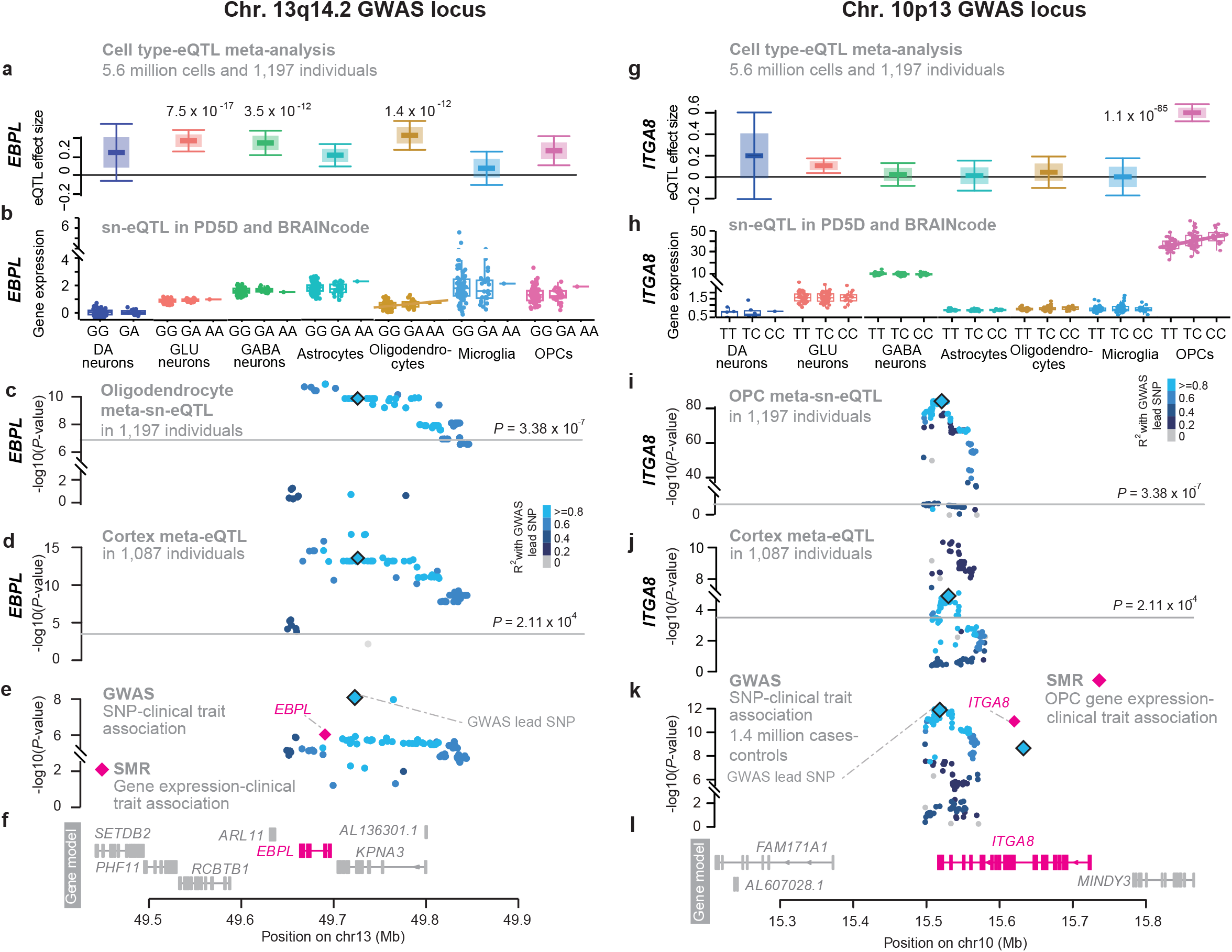
Functional map of the Chr. 10p13 and Chr. 13q14.2 loci. **a-f**, The Emopamil-Binding-Related Protein gene (*EBPL*) is a novel risk gene *cis*-regulated in oligodendrocytes as well as glutamate and GABA neurons. a, Effect sizes (beta coefficients) from cell type-eQTL meta-analysis across 1,197 individuals. **b**, genotype-expression associations in cell types from PD5D and BRAINcode. **c-d**, Locus-zoom plots show eSNP–EBPL associations **c**, in oligodendrocytes from 1,197 individuals and **d**, in cortex from another 1,182 individuals. **e**, SMR causality analysis shows a significant association between ***EBPL*** expression and PD risk (magenta diamond and font) overlayed on GWAS SNP-trait associations. **f**, seven genes are physically localized under this GWAS peak. **g-l**, The Integrin Subunit Alpha 8 gene (*ITGA8*) was preferentially expressed and specifically *cis*-regulated in OPCs --- implicating a novel cell type in the pathobiology of PD. **a**, Cell type-eQTL meta-analysis (*P* in OPCs = 1.1 × 10^-85^ using random effects model meta-analysis. **b**, Genotype-expression associations in cell types from PD5D and BRAINcode. **c-d**, Locus-zoom plots show eSNP–*ITGA8* associations in OPCs and **d**, in cortex from another 1,182 individuals. **e**, SMR causality analysis shows a significant association between ***ITGA8*** expression and PD risk overlayed on GWAS SNP-trait associations. **f**, four genes are physically localized under this GWAS peak. Box-and-whisker plots show the median (center line), the interquartile range (box), and the whiskers, which extend to 1.5x the interquartile range.

**Fig. 7.**
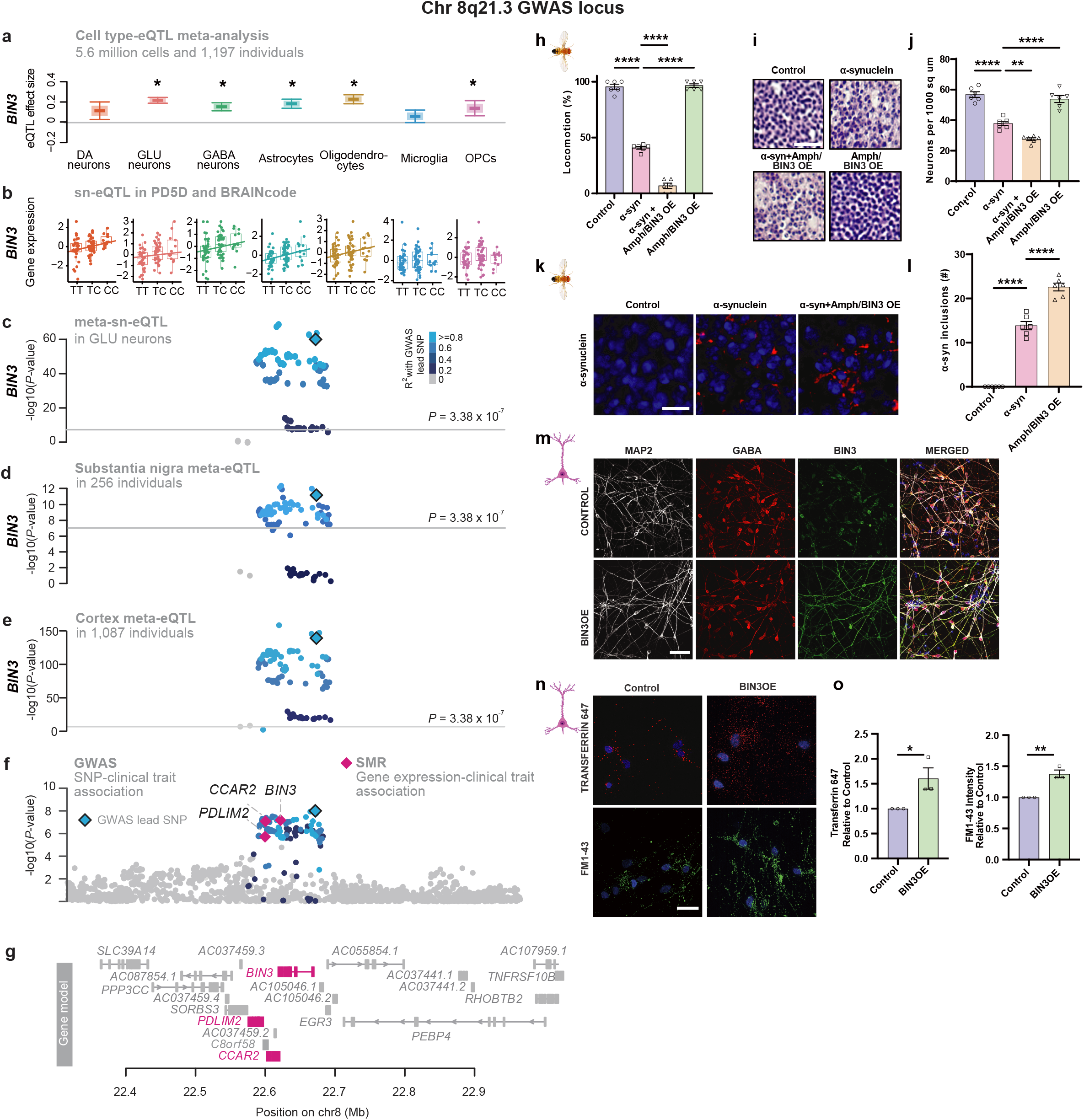
*BIN3* is a regulator of activity-dependent endocytosis and alpha-synuclein-induced neurodegeneration driving risk of PD. **a–g**, Noncoding variants at the Chr. 8q21 PD GWAS locus regulate *BIN3 e*xpression in neurons and glia by sn-eQTL. **a**, cell type-eQTL meta-analysis in 1,197 individuals. **b**, Genotype-expression associations in cell types from PD5D and BRAINcode. **c-e**, Locus-zoom plots show eSNP–*BIN3* associations (x-axis: chromosomal position; y-axis: *P* values), with R^2^ color-coding and the lead GWAS SNP marked by a black diamond. **c**, Cell type-eQTL results in glutamate neurons. **d**, The eQTL effect replicated in substantia nigra of 256 individuals and **e**, in cortex from 1,087 individuals. **f**, SMR causality analysis shows a significant association between increased *BIN3* expression and PD risk (magenta diamond); *PDLIM2* and *C8orf58* also shows association. Dots represent GWAS SNP–PD trait associations in the locus from Ref.^46^. **g**, Of 21 genes in the locus, sn-eQTL analysis identifies *BIN3* (magenta) as functional PD-linked target in glutamate and GABA neurons, astrocytes, and oligodendrocytes. *PDLIM2* and *C8orf58* (magenta) are additional target genes in this locus. Box-and-whisker plots show the median (center line), the interquartile range (box), and the whiskers, which extend to 1.5x the interquartile range. **h-o**, Overexpression of Amph/*BIN3* worsens alpha-synuclein toxicity in *Drosophila* and perturbs endocytosis in human PSC-derived neurons. **h**, Locomotion deficits in alpha-synuclein transgenic flies are exacerbated by Amph overexpression (N ≥ 60 per genotype). **i,j**, Hematoxylin staining shows increased neuronal degeneration with Amph overexpression (N = 6). **k,l**, Immunofluorescence shows increased alpha-synuclein aggregates in fly brains with Amph overexpression (N = 6). Mean ± SEM; one-way ANOVA with Bonferroni; ^**^P < 0.005, ^**^P < 0.0001; scale bars: 10 μm (i), 5 μm (k). **m-o**, *BIN3* overexpression enhances clathrin-mediated endocytosis in hPSC-derived neurons. **n**, Representative staining of *BIN3*-overexpressing and control GABAergic cultures. **n,o**, *BIN3* increases uptake of transferrin (KCl-treated) and FM1-43 after starvation, quantified in (o). Mean ± SEM; unpaired t-test; ^*^P ≤ 0.05, ^**^P ≤ 0.01; scale bars: 20 μm (m,n).

#### Neuronal endo-lysosomal dysfunction is an intense focus of investigation^64^

Key neuron-directed regulators included *STX1B* (vesicle fusion), *SH3GL2* and *BIN3* (membrane curvature)(**Fig. 7**), and *FLOT1* (clathrin-coating), *KTN1*, and *SCARB2* (lysosomal trafficking). *SCARB2* transports β-glucocerebrosidase, mutated in ∼10% of apparently sporadic PD cases. *NSF* and additional neuronal candidates (*CLCN3, SH2B1, ITGA2B, MAPK1IP1L, HSPA1B*) may also modulate vesicle cycling; *NSF* regulation may be sensitive to Lewy body burden in DA neurons (nominal dynamic-eQTL; **Supplementary Table S9**). *HSPA1B* emerged as a major chaperone hub (**Fig. 5**).

Taken together, PD GWAS variants converge on vesicle-cycle pathways across both glia and neurons, revealing a shared genetic architecture in which more than two dozen quantitative risk genes target vesicle biology, affecting myelin and neuroinflammation in glia and synaptic and endo-lysosomal dynamics in neurons.

#### Oligodendrocyte precursor cells (OPCs) as modulators of α-synuclein propagation

The quantitative risk genes *ITGA8* **(Fig. 6g-l)** and *GPNMB* implicated OPCs in controlling α-synuclein spread. We identified *ITGA8* as putative risk gene exclusively regulated in OPCs by the Chr. 10p13 locus. *ITGA8* modulates alpha-synuclein cell-to-cell transfer in cultured cells^65^. *GPNMB*, a facilitator of α-synuclein propagation^66^, showed PD-associated regulatory effects concentrated in OPCs and microglia, refining assumptions from bulk expression and consistent with recent studies^31-33^.

## Discussion

We define the genetic circuitry of Parkinson’s disease through one of the largest single-nucleus eQTL and bulk brain eQTL meta-analysis to date, spanning 2,279 human brains (**Table 1, Fig. 1**). Across nine cell types in cortex and substantia nigra, PD-associated GWAS loci cis-regulate 125 risk genes—62 of them novel—nearly doubling the number of putative causal genes and assigning genes and cell types to more than half of GWAS signals. Although PD has been framed as a neuronal alpha-synuclein disease^1,2^, 51% of functional risk genes are regulated in glia, with strong enrichment in oligodendrocytes (**Figs. 1,2,3,5**). These findings are consistent with prior evidence for oligodendrocyte heritability enrichment^24^ and white-matter microstructural alterations in PD^67^. Together, they position glial cell types—particularly oligodendroglia—as major mediators of common variant risk and suggest a mechanistic continuum between neuronal synucleinopathies and multiple system atrophy, a glial synucleinopathy centered on oligodendrocytes.

Across the 90 known PD GWAS loci, approximately 4,595 genes reside within associated intervals, limiting functional interpretation. Our single-cell framework resolves this by prioritizing 125 putative causal genes through convergent evidence from cell-type–specific eQTL mapping, colocalization, Mendelian randomization, and locus heterogeneity testing. We further establish a stringent causal inference framework by integrating allelic imbalance within single nuclei of heterozygotes (snASE) with orthogonal molecular and functional validation.

These results indicate that cell-type–specific gene regulation is a primary mechanism underlying common, sporadic PD. Moreover, *cis*-regulatory pleiotropy is frequent: 60% of resolved GWAS peaks harbored multiple eGenes, with an average of 2.5 cis-regulated targets per signal. This deviates from the classical one-locus-one-gene model and instead supports coordinated perturbation of gene networks as a key driver of PD risk^40^.

At the pathway level, PD risk converges on a glial–neuronal vesiculopathy network linking vesicle trafficking, antigen presentation, and α-synuclein biology. A central hub involves vesicle cycling and SNARE assembly (**Fig. 5**) with 25 risk genes partitioning into glial and neuronal programs. Neuronal risk genes regulated endocytosis, membrane curvature sensing, clathrin-mediated recycling, and lysosomal trafficking, extending established mechanisms in PD^64^. We identify BIN3, encoding a membrane-bending BAR domain protein required for vesicle formation, as the effector of the chr. 8q21.3 PD locus and show that it links perturbed vesicle cycling to α-synuclein toxicity in human neurons and a *Drosophila* model (**Fig. 7**).

In contrast, glial vesicle-cycle genes were enriched for myelin remodeling, endosomal trafficking, and neuroinflammatory secretion. The identification of *EBPL* (**Fig. 6**) and *FAM126A* as oligodendroglial PD risk genes is notable given their potential links to PI4P-dependent vesicle docking^60^ and myelin cholesterol metabolism^63^. OPC-specific regulation of *ITGA8* (**Fig. 6**) and glial enrichment of *GPNMB* refine earlier bulk-tissue interpretations^65,66^. Both genes modulate alpha-synuclein transmission *in vitro*, suggesting that OPCs may influence cell-to-cell propagation of alpha-synuclein during disease progression.

Twenty-seven quantitative risk genes mapped to the linked antigen-presentation pathway (**Fig. 5**), including 17 forming an interconnected protein–protein interaction network hub. We identified five HLA genes associated with PD risk in glia and cell-type–specific regulation of MHC loading machinery such as *TAP2* and *ABCB9*. Notably, *ABCB9* regulation was exclusive to oligodendrocytes, implicating oligodendroglial antigen processing as a previously unrecognized PD mechanism.

We also observed convergence between and common and rare genetic forms of PD within the glial-neuronal vesiculopathy network (**Fig. 5i**). Quantitative risk genes showed direct protein–protein interactions with monogenic PD genes, including DNAJC6, *SYNJ1, LRRK2*, and *SNCA*, supporting shared pathogenic pathways across familial and sporadic disease.

This study has limitations. Gene expression does not fully predict protein abundance^68^, and methodological differences between dopaminergic and cortical cell-type analyses complicate direct comparisons. While our data support gene regulation as a primary mechanism of common PD risk, other mechanisms— including coding variation at loci such as *SNCA, GBA, LRRK2*, and *TMEM175*^69,70^—also contribute. Ambient RNA could influence expression^71^, but its removal^71^ had no material effect on expression of the risk genes (R ≥ 0.994 with vs. without removal). Larger sample sizes, integration of splicing and additional molecular modalities, and extension to other vulnerable cell types will further resolve remaining GWAS signals. Proxy-case bias has been found in AD GWAS^72^, but does not affect PD lead GWAS variants^46^, and sensitivity analyses revealed no evidence of downstream bias in our study, consistent with prior work^46^.

In conclusion, this work provides a cell-type–resolved atlas of gene-regulatory mechanisms underlying PD, linking common genetic variation to specific effector genes, pathways, and cell types. It supports a model in which PD is, to a substantial extent, a gene-regulatory disorder involving both glia and neurons, with oligodendroglia playing a central and previously underappreciated role. By systematically bridging GWAS signals to their cellular and molecular targets, this framework enables mechanistic follow-up, therapeutic target prioritization, and future precision medicine strategies for PD.

## Materials and Methods

### Sample collection and processing methods for sn-eQTL analyses

#### Brain samples

In our Parkinson’s disease in 5D (PD5D) cell atlas project, postmortem brain samples were obtained from the Banner Sun Health Research Institute Brain and Body Donation Program in Sun City, Arizona^73^. All subjects had annual movement and cognitive evaluations prior to death and a complete neuropathologic examination as described previously^73^, which includes the Unified Staging System for Lewy Body Disorders^74^. All brain samples were obtained through a rapid autopsy program with superb post-mortem intervals (median PMI, 3 hours) and high RNA integrity numbers (median RIN, 9.2; indicating excellent RNA quality). Detailed quality measures and demographic characteristics of these high-quality, frozen postmortem samples are shown in **Supplementary Table S1**.

Our study included brains from participants spanning the entire disease process from healthy controls to incipient, prodromal PD-associated Lewy body neuropathology (“incidental Lewy body disease”) to clinically manifest Parkinson’s disease in order to dynamically capture and exam the disease context on gene expression. Age- and sex-similar individuals in each of the three diagnostic groups of healthy controls (*N* = 30); prodromal, PD-associated Lewy body neuropathology (*N* = 29); and PD (*N* = 35) were included.

Cases with prodromal, PD-associated Lewy body neuropathology are those not meeting clinical diagnostic criteria for PD or other neurodegenerative diseases but found with Lewy body inclusion at autopsy. These cases are widely considered a preclinical stage of PD ^75^, providing an opportunity to investigate the dynamic eQTL effects of genotypes and Lewy body neuropathology on gene expression. PD cases had a clinical and neuropathological diagnosis of PD.

Healthy control subjects were defined with the following stringent inclusion and exclusion criteria. Inclusion criteria: (1) absence of clinical or neuropathological diagnosis of a neurodegenerative disease e.g., Parkinson’s disease according to the UKPDBB criteria ^76^, Alzheimer’s disease according to NIA-Reagan criteria ^77^, dementia with Lewy bodies by revised consensus criteria ^78^. (2) PMI ≤ 48 hours; (3) RIN ^79^ ≥ 6.0 by Agilent Bioanalyzer (good RNA integrity); (4) visible ribosomal peaks on the electropherogram.

Exclusion criteria for all autopsy brains were: (1) a primary intracerebral event as the cause of death; (2) brain tumor (except incidental meningiomas); (3) systemic disorders likely to cause chronic brain damage.

Our research complies with all relevant ethical regulations. Sample collection was approved by the Institutional Review Board of Banner Health. The study was approved by the Institutional Review Boards of MassGeneral Brigham and of Yale University.

#### Nuclei isolation and library preparation

Nuclei were isolated from frozen human temporal cortex tissue using a previously described protocol (Habib et al 2017; pmid-28846088) with modifications (Protocol.io for more details). Total of 10 sections (50 µm thick) of human temporal cortex tissues were homogenized in ice-cold lysis buffer (LB) (prepared using Nuclei PURE lysis buffer (Sigma, #NUC201) plus DTT (1:1000; DL-Dithiothreitol, Sigma) plus Triton X-100 (0.01%, Sigma)) using a pestle (Sigma #D8938-1SET). Tissue homogenates were incubated on ice for 5min, filter through a 70 µm cell strainer (Thermo Fisher, #2236548) and then centrifuged at 500x g, 4°C for 5min. The supernatant was removed, and the pellet was resuspended in LB, then incubated on ice for 5 min followed by centrifugation at 500x g for 5 min at 4°C. The nuclei pellets were subsequently washed with Nuclei Wash and Resuspension Buffer (NWRB), prepared with 0.01% BSA (Thermo Fisher) and freshly added RNase inhibitors (40 U/ml, Clontech; #2313A) in 1X DPBS (Gibco, #14190-144). Next, the nuclei were centrifuged at 500x g for 5 min at 4°C, and the washing step was repeated once more. Isolated nuclei pellet was resuspended in NWRB containing DAPI (0.01 mg/ml; Thermo Fisher, #D1306) and filtered through the 40 µm Flowmi cell strainer (Bel-art, #H13680-0040). Next, DAPI-stained nuclei were sorted using FACSAria^™^ III cell sorter (BD Biosciences) equipped with required lasers and a 100 µM nozzle. DAPI-positive nuclei were detected using a violet laser (405 nm) with a 450/50 bandpass filter. Before sorting, appropriate gating parameters were employed: nuclei were first gated based on their forward versus side scatter properties (FSC-A at 5V / SSC-A at 180V) and then gated based on their DAPI expression. Approximately 90,000 events were collected in 1.5 mL Eppendorf tubes and counted on a hemocytometer for quality assurance. Next, 30,000 -40,000 events (∼15,000 nuclei) per sample were used for 10x Genomics Chromium loading. The Gel Beads-in-emulsion (GEM) generation and barcoding was performed using Chromium Next GEM Single Cell 3 ′ Gel Bead Kit v3.1 (PN-1000122 and PN-1000123) and 10x Chromium Controller (10x Genomics) as per the manufacturers protocol. Indexed sequencing libraries were constructed using the Single-Cell 3′ Library Construction Kit v3.1 (10x Genomics, #PN-1000190) and Dual index kit TT set A (10x Genomics, #PN-1000215), as per the manufacturer’s protocol. The barcoded single nuclei libraries were quantified by quantitative PCR using KAPA Library Quantification Kit (KAPA Biosystems/Roche, #7960336001 and kit code-KK4873). Next, the libraries were sequenced using NextSeq 500/550 High Output Kit v2.5 150 Cycles (Illumina, #20024907) on an Illumina NextSeq 550 platform with following run parameters: Read 1: 28; i7 index: 10; i5 index: 10; Read2: 120.

#### Genotyping and processing

All subjects were genotyped by Multi-Ethnic Genotyping Array (MEGA) using the Infinium Multi-Ethnic Global-8 v1 kit (Illumina, #WG-316-1001) following manufacturer’s instructions. We applied PLINK (v1.9beta) and in house scripts to perform rigorous subject and SNP quality control (QC) for each dataset in the following order: (1) Set heterozygous haploid genotypes as missing for all chromosomes, including autosomes and chromosome X; (2) remove variants with GenCall score (GC) < 0.15; (3) remove variants which could not be mapped to GRCh38/hg38 reference genome; (4) remove subjects with call rate < 95%; (5) remove subjects with sex discordance; (6) remove SNPs with genotype call rate < 95%; (7) remove SNPs with Hardy-Weinberg Equilibrium testing P value < 1×10-6; (8) remove SNPs with informative missingness test (Test-mishap) P value < 1 ×10-9; (9) remove SNPs with minor allele frequency (MAF) < 0.01; (10) remove subjects with outlying heterozygosity rate based on heterozygosity F score (beyond 4-fold sd from the mean F score); (11) IBS/IBD filtering: pairwise identity-by-state probabilities were computed for removing both individuals in each pair with IBD>0.9 and one subject of each pair with IBD > 0.1875; (12) Subjects were integrated with 1000 Genomes Project phase3 individuals and tested for population substructure by performing principal components analysis (PCA) using PLINK. Based on first and second principal components, significant genotypic outliers (non-European individuals) were excluded.

#### Imputation of genotypes

After quality control, genotype data of all subjects were combined with 4,020 European individuals reported in our previous work ^80^ and were submitted to the TOPMed Imputation Server for genotype imputation. During the imputation, TOPMed r2 reference panel which includes 97,256 reference samples and 308,107,085 genetic variants distributed across the 22 autosomes and the X chromosome was used as the reference panel, Eagle(v2.4) and Minimac4 were selected for phasing and imputing, respectively. After imputation, variants with R2 < 0.3 or MAF < 0.001 were excluded from further analysis. The correlation between real MEGA-chip genotype and the imputed genotype was employed for evaluating the accuracy of the imputed genotypes.

### Single-cell eQTL analysis of the Parkinson Cell Atlas in 5D (PD5D)

#### Single-cell data preprocessing and quality control

The Cell Ranger Single Cell Software Suite (v5.0.1) provided by 10x Genomics was used to proceed and raw sequencing data into FASTQ files and demultiplex the data into separate pools of samples as well. FastQC (v0.11.9) and MultiQC (v1.9) were used to assess base quality, then the Cell Ranger count function (--include-introns) was employed for aligning the reads of each sample to GRC38/hg38 reference transcriptome, and investigate the raw UMI counts of genes on the basis of gene annotation for GRC38/hg38 reference transcriptome from the Cell Ranger website (refdata-gex-GRCh38-2020-A_Ensembl98). Quality control was conducted to remove low quality cells, samples and lowly expressed genes. Cells having >5% of reads originating from the mitochondria, < 200 unique genes or whose number of unique genes > three times of median absolute deviation of number of unique genes across all cells (9,378 UMIs) were removed by using Seurat (v4). Genes expressing in ≤ 3 cells of each individual were set to no expression. After the cell and gene level quality control, sample level bulk expression of all genes were calculated by in-house script (https://zenodo.org/records/17611730). Principal components analysis, hierarchical clustering and relative log expression (RLE) plots towards the sample level bulk gene expression matrix were then employed to evaluate sample quality and detect outlier samples. Samples which are outliers in all of the analysis (PCA, hierarchical clustering and RLE) and have less than 2,000 cells were removed from the analysis.

#### Normalization, dimensionality reduction, covariates correction and cell type clustering

The aggregated matrix was then processed using a Seurat (v4.3.0) workflow in RStudio (v4.0.2). The count data was normalized to give counts per ten thousand and these values + 1 were then natural log transformed (NormalizeData, params: normalization.method = “LogNormalize”, scale.factor = 10000). Next, the 2000 most variable genes were identified (FindVariableFeatures, params: selection.method = “vst”) and the data scaled (ScaleData) across all genes. The 2000 most variable features were used for dimensionality reduction using glmPCA (v0.2.0) (RunGLMPCA). Harmony (v1.2.0) was run to account for covariates, correcting the glmPCA for the scRNA-sequencing batch, sample, age bracket (binned into 10 year intervals), RIN bracket, PMI bracket and sex, with theta being set at 0.4 for each covariate. An elbow plot was used to help determine the ideal number of harmony dimensions (60) to pass to the FindNeighbours function, which was followed by cluster identification using the Leiden algorithm, in which a range of resolutions was tested (0.5 – 1.6), in order to identify a suitable value (FindClusters, params: resolution = 1.5, algorithm = 4, method = “igraph”). UMAP was then run (RunUMAP) to visualize the resulting clusters. Clusters with less than 200 cells or representing less than 20 samples were removed as part of cluster quality control. Doublet cells were identified using the scDblFinder package (v1.12.0), by running the function scDblFinder on a per sample basis. All doublet cells were removed as well as any clusters that consisted of 30% or more doublet cells. The UMAP was subsequently re-run to clean up the visualization. The cell types were annotated by using canonical marker genes for brain cells, including SLC17A7 and SLC17A6 for GLU neurons, GAD1 and GAD2 for GABA neurons, AQP4 and GFAP for Astrocytes, PLP1 and MOBP for Oligodendrocytes, CX3CR1 and P2RY12 for Microglia, FLT1 and ABCB1 for Endothelial cells, VCAN and BCAN for OPCs and PDGFRB for Pericytes.

#### PD risk variants selection

To exploit potential expression regulatory effect of 90 GWAS loci revealed by the genome-wide investigation of 26,035 PD cases and 403,190 controls ^46^, three types of risk variants were selected for ensuring the inclusive of potential causal expression regulatory variants ^81^: 1. PD GWAS significant variants which have a GWAS p-value ≤ 5 × 10^-8^; 2. PD GWAS sub-threshold variants which have a GWAS p-value ≤ 1 × 10^-6^ and locate around 2Mb of PD GWAS loci; 3. Proxy variants which have linkage (R2 ≥ 0.4) with either PD GWAS significant variants or PD GWAS sub-threshold variants in 1000 Genomes phase3 EUR population genotype data. Variants which have minor allele frequency (MAF) < 0.05 in our genotype data or without having positional information in GRC38/hg19 reference genome were excluded from the further analysis. In total, 32,267 variants, including 7,057 PD GWAS significant variants, 3,550 PD GWAS sub-threshold variants and 21,660 proxy variants of PD GWAS significant variants or PD GWAS sub-threshold variants, were selected as the candidate variants in our analysis.

#### Single-nucleus cis-eQTL (sn-eQTL) mapping

For each cell, gene expression was normalized by dividing the gene’s UMI count by the total UMI count in that cell and multiplying by a scaling factor of 10,000. For each cell type, pseudo-bulk subject expression for each gene was calculated by summing the gene’s expression values across all cells of that cell type from the same subject and dividing by the number of those cells. This is equivalent to the average expression level of the gene across all cells of that cell type for that subject. These pseudo-bulked expression values were then used as input for the cell type– specific eQTL analysis.

Surrogate variable analysis (SVA) and ComBat packages were used in R^82^. ComBat^82^ was used to adjust for sequencing batches. ComBat applies an empirical Bayes framework to adjust for batch effects in both the mean (e.g., a batch with consistently higher expression) and the variance (e.g., a batch with greater variability) of gene expression across samples. Gene expression values adjusted with ComBat were then used as input for surrogate variable analysis. Hidden confounders were estimated through surrogate variable analysis. A linear model implemented in the fsva function in the SVA package (v3.38.0) was used to adjust for the known covariates of age, sex, RNA integrity number, and post-mortem interval as well as hidden covariates (the first 30 SVA low-dimension embedding factors).

The resulting covariate-adjusted gene expression values were normalized to z-scores by subtracting the mean from each value and dividing by the standard deviation using the R function scale, which forces the expression values to follow a standardized normal distribution (with a mean of 0 and a standard deviation of 1 across individuals). That makes the unit of β = “standard deviations of expression per allele.” PD-associated risk variants located in a genic region or 1 Mb around the gene were used for mapping single-nucleus cis-eQTL of that gene in each cell type with the R Package Matrix EQTL (v.2.3).

#### Cell type-eQTL meta-analyses

Meta-analysis across the three single-nucleus eQTL datasets, including the single-nucleus eQTLs from PD5D (this study), from Fujima et al.^32^, and from Haglund et al. ^31^, was conducted in each of the eight cortical cell types to increase power by using Metasoft (https://hanlab-snu.github.io/METASOFT/). For dopamine neurons, dopamine neuron eQTLs in BRAINcode^40^ were mapped using the same pipeline as the single cell eQTL analyses described above, and then the meta-analysis was conducted by using the dopamine neuron eQTLs from BRAINcode and the eQTLs from 52 days cultured dopamine neurons derived from human pluripotent stem cells^41^. For each variant–gene pair, the effect size and its estimated standard error from each eQTL dataset was used as input for meta-analysis using Metasoft to improve the power of mapping sn-eQTLs. Three different models were conducted during the meta eQTL analyses, including the Fixed effects model (FE model) analysis was based on inverse-variance-weighted effect sizes, the Random effects model (RE model) analysis was a conservative model based on inverse-variance-weighted effect sizes, and the Han and Eskin’s random effects model (RE2 model) was optimized to detect associations under heterogeneity. P-values from the RE2 model and the effect sizes and the standard errors of effect sizes from the RE model were used in the following analyses. sn-eQTL nominated *P* ≤ the Bonferroni-threshold of 3.38 × 10^-7^ (e.g, 0.05/147,929 independent eSNP-eGene pairs) was employed as the cutoff to identify significant sn-eQTLs in each cell type. LD-clumping was conducted to identify independent eQTLs of each significant eGene using the PLINK (v1.90) as following command with the genotypes of 1000 Genome Projects EUR population as the LD reference: --clump eSNPs --clump-p1 3.38 × 10^-7^ --clump-p2 3.38 × 10^-7^ --clump-r2 0.1 --clump-kb 500. In this scenario, each independent sn-eQTL of eGenes was a sn-eQTL LD chunk which is LD-independent from other sn-eQTLs(or sn-eQTL LD chunks). sn-eQTLs from different cell types and locating on the same LD chunk was thought as the same sn-eQTL. Cell types of each of the independent sn-eQTLs were exploited as following: any eSNP from a sn-eQTL LD chunk passed the bonferroni-threshold of 3.38 × 10^-7^ in a cell type indicated the sn-eQTL was detected in that cell type. To explore potential inflation of the cell type preferential sn-eQTLs in other cell types, we extracted the p-values of eSNPs of the sn-eQTLs in other cell types and employed the π_1_ statistic to estimate the proportion of eSNPs whose p-values in other cell types follow non null distribution. The above-described code can be found here (https://zenodo.org/records/17611730).

#### Colocalization analysis

GWAS summary statistics from Ref. ^25^ and eQTL summary statistics for all genes for each cell type were used for co-localization analysis using coloc (v5.2.3) REF. LD clumping was conducted to identify independent GWAS LD chunks of each locus using the PLINK (v1.90) similar to what is described above using genotypes of the EUR population from the 1000 Genomes Project as the LD reference: --clump eSNPs --clump-p1 5.0 × 10^-8^ --clump-p2 5.0 × 10^-8^ --clump-r2 0.1 --clump-kb 500.

For each cell type, genes with sn-eQTL associations passing the *P* ≤the Bonferroni-threshold of 3.38 ×10^-7^ were used for the colocalization analysis. The priority of colocalization was set as following: prior probability of a SNP-trait association (PP1) = 1e^-4,^ prior probability of SNP-gene association (PP2) = 1e^-4^; PP that a variant encodes affects both traits PP12 = 1e^-5^. Nominal p-value, effect size, standard error of the effect size, sample size and minor allele frequency of eQTLs and GWAS signals were employed during the colocalization analysis. Single causal variant assumption was applied for colocalization analysis, as the PD GWAS locus had been splitted into independent GWAS LD chunks. A posterior probability of ≥0.8 (PP4 in the result) was used as cutoff for strong GWAS-eQTL colocalization.

#### Summary based Mendelian randomization (SMR)

GWAS summary statistics for Parkinson’s disease (PD) risk variants from from Ref. ^25^ were used for SMR analysis. In each cell type, eGenes with significant eQTLs (e.g, cell type eQTL p-values ≤ 3.38 × 10^-7^) were forwarded for SMR analysis. Before conducting the analysis, effect alleles and direction of effect were aligned to the GWAS summary statistics. PD GWAS risk variants located within 1 Mb of a significant eGene were used for the SMR analysis using the SMR (v1.3.1) tool. The sn-eQTL summary statistics of these eGenes, including beta coefficient, standard error of the beta coefficient, nominal meta-eQTL *P*-values, alleles, and effect allele frequency from the 1000 Genomes Project phase3 EUR population, were employed for SMR analysis. These sn-eQTL summary statistics were converted into a binary format suitable for SMR with the SMR tool. During the SMR analysis, the 1000 Genomes Project phase3 EUR population was used as reference to identify linkage among the variants. Considering the complexity of PD GWAS loci which may harbor multiple signals in the same locus, both single-causal variant and multiple-causal variants SMR tests were conducted. *P*-values from single-causal variant SMR tests or multiple-causal variants SMR tests equal or lower (more significant) than the Bonferroni threshold for significance of 1.18 × 10^-4^ (e.g. 0.05/271 forwarded eGenes) were considered evidence for associations between gene expression and PD risk. Using all eSNPs for the eGenes maximizes the power of the SMR test.

#### HEIDI test for linkage

Subsequently, the HEIDI test was implemented to distinguish pleiotropy from linkage between eSNPs and PD GWAS risk variants for each of the eQTL chunk and PD GWAS signal chunk pairs, which were assumed to harbor a single causal variant each for each the eQTL and the PD GWAS signal. The eGenes which passed the HEIDI test (HEIDI test *P*-value ≥ 0.01) in the cell type which they were significantly associated with PD were considered a candidate risk genes for further analyses.

### Characterization of putative causal risk genes

#### Single nucleus Allele-specific expression (ASE) analysis

We forwarded significant sn-eQTLs for snASE analysis as we reported recently in Ref.^83^. We performed snASE analysis by using a Nextflow V2^84^ pipeline. In brief, the pipeline takes in fastq files from 10X and a phased vcf. We mapped snRNA-seq reads against the reference genome by using the STARSolo pipeline^85^, running STAR with WASP^86^ capability to help mitigate the effects of reference bias. More specifically, we ran STAR with the arguments described in https://github.com/seanken/ASE_pipeline/blob/main/QuantPipeline.no.conda.nfline103. The resulting bam file was then processed using a htsjdk ^87^ (version 2.21.1) based Java application. More specifically, htsjdk was used to read each read into Java, and for each read the UMI, CBC, Gene information, and allele (alternative *vs*. reference) were extracted from the associated tags in the bam file (for the allele, we used the information in the vA tag), excluding multimapped reads and those that did not pass the WASP filter. This information was then added to a HashMap, which was later used to assign UMI/CBC/gene triplets to an allele and produce a cell level ASE matrix. This data was loaded into R with the scAlleleExpression package which was also built by our previous study^83^, using the GetSNPs command, resulting in a pseudobulk ASE dataframe (pseudobulked by sample of origin and cell type). We tested for ASE difference between alleles by fitting a betabinomial generalized linear model on this pseudobulked ASE data for each cell type, where fitting the betabinomial model was done with the TestSNP_aod command in scAlleleExpression, which works by fitting a betabinomial model with the betabin command in the aod package^88^. Only variants with ≥ 20 phased UMIs were retained in the study. The snASE analysis was employed for validating sn-eQTLs purpose and we used a *P* value ≤ 0.05 as a cutoff for estimating snASE significance.

### Evaluation of eQTLs across seven human cortex data sets and two substantia nigra data sets

#### Cortex and substantia nigra eQTL methods

Seven cortex cohorts and two substantia nigra cohorts were analyzed as we previously described in Ref.^36^. Five of these datasets used RNA-sequencing technology for estimating gene expression, two used gene expression microarrays. Four used whole genome sequencing (WGS) for genotyping; three used genotyping arrays as we detailed in Ref.^36^. All datasets were accessed under Data Use Agreements and local IRB approval.

#### Data sets used for human cortex and substantia nigra expression Quantitative Trait Locus (eQTL) analysis

Human cortex eQTL analysis was performed using seven bulk cortex data sets, and human substantia nigra eQTL analysis was performed using two bulk substantia nigra data sets (Table S4). Only the transcriptome data from the cortex/substantia nigra of healthy controls of each cohort were included in the study (Table S4). ROSMAP, MayoRNAseq, MSBB, and HBTRC data (bulk cortex) were obtained from the AD Knowledge Portal (https://adknowledgeportal.org) on the Synapse platform (Synapse ID: syn9702085). CommonMind (bulk cortex) was obtained from the CommonMind Consortium Knowledge Portal (https://doi.org/10.7303/syn2759792) also on the Synapse platform (Synapse ID: syn2759792); GTEx (bulk cortex + bulk substantia nigra) was obtained from https://gtexportal.org/home/. UKBEC (bulk cortex and bulk substantia nigra), was obtained from http://www.braineac.org/. The data sets are described in detail at each of the source portals and in the corresponding original publications.

#### Bulk cortex gene expression data and bulk substantia nigra gene expression processing for eQTL analyses

For RNAseq datasets, using RStudio (v4.0.2) gene reads count was normalized to TPM (Transcripts Per Kilobase Million) by scaling gene length first and followed by scaling sequencing depth. The gene length was considered the union of exon length. Consistent and stringent quality control and normalization steps were applied for each of the cohorts: (1) For sample quality control, samples with poor alignment were removed. Samples with > 10 M mapped reads and > 70% mappability by considering reads with mapping quality of Q20 or higher (the estimated read alignment error rate was 0.01 or less) were kept. (2) Removing sample mix-ups by comparing the reported sex with the transcriptional sex determined by expression of the female-specific *XIST* gene and the male-specific *RPS4Y1* gene. (3) Removal of sample outliers: Sample outliers with problematic gene expression profiles were detected based on Relative Log Expression (RLE) analysis, Spearman correlation based hierarchical clustering, D-statistics analysis ^89^. (4) Normalization: Gene expression values were quantile normalized after log10 transformation by adding a pseudocount of 10^-4^. (5) Batch effects were removed with sva (v3.38.0) using the combat function and adjusting for age, sex, RIN, PMI and latent covariates using the fsva function. Residuals were outputted for downstream analyses. For array-based gene expression datasets, directly downloaded, quality-controlled gene expression profiles were used.

#### Genotype data processing for bulk cortex and bulk substantia nigra eQTL analyses

The PLINK (v1.9beta) package and in-house scripts (10.5281/zenodo.13333397) were used to perform rigorous subject and SNP quality control (QC) for each dataset in the following order: (1) Set heterozygous haploid genotypes as missing; (2) remove subjects with call rate < 95%; (3) remove subjects with sex discordance; (4) remove SNPs with genotype call rate < 95%; (5) remove SNPs with Hardy-Weinberg Equilibrium testing P value < 1 × 10^-6^; (6) remove SNPs with informative missingness test (Test-mishap) P value < 1 × 10^-9^; (7) remove SNPs with minor allele frequency (MAF) < 0.05; (8) remove subjects with an outlying heterozygosity rate based on the heterozygosity F score (more than 4*sd from the mean F score); (9) IBS/IBD filtering: pairwise identity-by-state probabilities were computed for removing both individuals in each pair for IBD > 0.98 and one subject of each pair for IBD > 0.1875; (10) population substructure was tested by performing principal components analysis (PCA) using smartPCA in EIGENSOFT ^90^. After exclusion of PCA outliers, the top 3 principal components were used as covariates for adjusting population substructures. Next, all genotype data were submitted for imputation using the TOPMed imputation server with the same tools and parameters as described previously for the genotyping of single-nucleus transcriptome samples.

#### Bulk cortex and bulk substantia nigra eQTL analysis

eQTL mapping in each cohort was conducted using the R Package Matrix EQTL (v2.3) by applying the additive linear model on a high-performance cluster. For cis-eQTL analysis, SNPs were included if their positions were within 1 Mb of the transcription start site of a gene.

#### Bulk cortex and bulk substantia nigra eQTL meta-analysis

Meta-eQTL analysis was performed using three separate effects models implemented in METASOFT (v2.0.1)^91^ using effect size and standard error of SNP-gene pairs from each dataset as input. Fixed effects model (FE model) analysis was based on inverse-variance-weighted effect sizes. Random effects model (RE model) analysis was a conservative model based on inverse-variance-weighted effect sizes. Han and Eskin’s random effects model (RE2 model) was optimized to detect associations under heterogeneity. The statistics resulting from the RE and RE2 model were employed in the following analyses. sn-eQTLs whose lead eSNPs which passed the cutoff of false discovery rate (FDR) ≤ 0.05 (e.g, the nominated p-value from RE2 model ≤ 2.11 × 10^-4^) were thought as successfully replicated in bulk cortical eQTLs or bulk substantia nigra eQTLs.

#### Gene set and disease trait enrichment analyses

Metascape^92^ was used to delineate gene sets significantly enriched in our list of 125 putative causal risk genes. For 116 of the 125 risk genes annotations were available and were used for enrichment analyses. Analogous down-stream analyses were performed for subsets of these risk genes regulated across glia cells, across neurons, and across each of the nine cell types. The remaining genes, generally non-coding genes, lacked annotations and were excluded from these analyses. For each given gene list, pathway and process enrichment analysis were carried using standard Metascape parameters, ontology sources, and methods as reported in Ref.^92^: KEGG Pathway, GO Biological Processes, Reactome Gene Sets, Canonical Pathways, CORUM, WikiPathways, and PANTHER Pathway^92^. All genes in the genome were used as the enrichment background^92^. Terms with a p-value < 0.01, a minimum count of 3, and an enrichment factor > 1.5 (the enrichment factor is the ratio between the observed counts and the counts expected by chance) were collected and grouped into clusters based on their membership similarities^92^. More specifically, P-values were calculated based on the hypergeometric distribution^92^. Kappa scores^92^ were used as the similarity metric when performing hierarchical clustering on the enriched terms, and sub-trees with a similarity of > 0.3 are considered a cluster. The most statistically significant term within a cluster is chosen to represent the cluster.

Enrichment analysis for disease traits was performed in DisGeNET^93^ through the Metascape interface. DisGeNET id one of the largest available collections of genes and variants associated with human diseases, integrating data from expert curated repositories, GWAS catalogues, animal models and the scientific literature.

#### Protein-protein network interaction analyses

were performed in Metascape^92^. For each risk gene list, protein-protein interaction enrichment analysis was performed with the following databases: STRING ^94^, BioGrid^95^, and Ref.^96^. Only physical interactions in STRING (physical score > 0.132) and BioGrid were used with Metascape default filters. The resultant network contains the subset of proteins that form physical interactions with at least one other member in the list. GO enrichment analysis was applied to the network. The network layout was visualized in Cytoscape^97^. Each term is represented by a circle node, where its size is proportional to the number of input genes fall under that term. The color represent its cluster identity (i.e., nodes of the same color belong to the same biological cluster from Metascape with ad-hoc updates based on expert review of the literature). Terms with a similarity score > 0.3 are linked by an edge (the thickness of the edge represents the similarity score).

### *Drosophila* experiments

#### *Drosophila* genetics

*Drosophila* crosses were performed at 25°C. Flies were aged at 25°C for 7 days for locomotion assays and 10 days for histological analysis. Equal numbers of male and female flies were used in each experiment. The pan-neuronal drivers *nSyb-GAL4* and *nSyb-QF2* were used for all experiments. We have previously described *QUAS-wild type α-synuclein* transgenic flies ^53^. The *UAS-Amph* stock (#6499) was obtained from the Bloomington *Drosophila* Stock Center.

#### Locomotion assay

Adult flies were aged in vials containing 9–14 flies per vial. At day 7 flies were transferred to a clean vial without anesthesia and the locomotion assay performed as described in detail ^98^. In brief, the vial was gently tapped three times to trigger the startle-induced locomotion response, then placed on its side for 15 seconds. The percentage of flies still in motion was then recorded.

#### Histology, immunostaining and imaging

*Drosophila* heads were fixed in formalin and embedded in paraffin before sectioning. Serial frontal sections (2 or 4 μm) of the entire brain were prepared and mounted on glass slides. To assess neuronal density, sections were stained with hematoxylin and imaged using the imaging software SPOT. The number of anterior medulla neurons in an approximately 1000 μm^2^ area were counted using the cell counter plugin in ImageJ. For immunostaining on paraffin sections, antigen retrieval was performed by microwaving slides in sodium citrate buffer for 15 minutes. Slides were then blocked in 2% milk in PBS with 0.3% Triton X-100, followed by overnight incubation with an antibody recognizing aggregated alpha-synuclein (5G4, Millipore, 1:1,000,000) and then with a secondary antibody coupled to Alexa Fluor 555. Imaging was performed with Zeiss laser-scanning confocal microscopy. The number of inclusions per 60 μm^2^ area was determined.

### iPSC experiments

#### Generation of GABA interneurons

These studies were approved by the stem cell research oversight committee, University of Wisconsin-Madison. Human embryonic stem cells (H9 or WA09, WiCell), were used to generate BIN3 overexpression (BIN3OE) stem cell line using CRISPR/Cas9 gene editing technology. Stem cells were grown on Matrigel (BD Biosciences, #354277) with TeSR-8 medium (Stemcell Technologies, #05990) to 40% confluency. For GABA interneuron differentiation, we used our previously published protocol^99^. Stem cells were treated with neural differentiation medium (DMEM/F12,#11330 + 1× N2 Supplement, #17502048 + 1 1x Glutamax, #35050 + B27 without Vitamin A, #1258701, all Gibco) with the SMAD inhibitors SB431542 (Stemgent, #040010), DMH-1 (Tocris Bioscience, #4126) (both at 2 μM) for 8 days. Media was changed every other day. On Day 8, neuroepithelial cells were lifted using Accutase and were patterned to GABA interneuron progenitors with the smoothened antagonist -SAG (Calbiochem, #566660, 1 μM) and FGF2 (R&D, #233-FB, 10 ng/ml) for 14 days. On Day 21, neural progenitors were dissociated with Accutase to single cells and plated on Poly D Lysine and Laminin coated coverslips in the maturation media (DMEM/F12, 1× B27 Supplement, 1× non-essential amino acids, 1× Glutamax) supplemented with BDNF (Peprotech, #450-02), GDNF (Peprotech, #450-10), cAMP (Sigma Aldrich, #D-0260), Ascorbic Acid (Sigma Aldrich, #A4403) and Compound E (0.1 μM, TOCRIS, #565790-500UG).

#### Transferrin Uptake assay

Human transferrin conjugated with Alexa Fluor 647 (Thermofisher, #T23366) was used for the endocytic uptake experiments. On Day 21 after maturation, neurons were briefly starved in basal media for one hour. Neurons were treated with Human Transferrin at 10 ug/ml for one minute at 37°C followed by fixation with 4% PFA(Sigma Aldrich, #P6148). Cells were washed three times with PBS and stained with Hoechst (Thermofischer Scientific, #R37165) for 10 minutes. After washing with PBS, coverslips were mounted using Fluoromount G (Thermofischer Scientific, #00-4958-02). Imaging was performed using Elyra 7 microscope (Zeiss). Images were captured using 40x objective. Statistical analysis was performed using Graphpad Prism 9.

#### FM1-43 Uptake assay

FM1-43 FX (Thermofisher Scientific, F35355) was used for the endocytosis experiments. On Day 21 after maturation, neurons were transferred to standard artificial cerebrospinal fluid (ACSF) containing 140 NaCl, 4 KCl, 2 CaCl_2_, 2 MgCl_2_, 10 HEPES, 5 D-glucose, pH 7.4 (adjusted with NaOH). Neurons were treated with 10 µM of FM1-43 FX for 10 minutes at room temperature. Then neurons were washed three times with ACSF without external calcium. After washing, neurons were fixed with 4% PFA for 20 minutes at room temperature. Cells were washed three times with PBS and stained with Hoechst for 10 minutes. After washing with PBS, coverslips were mounted using Fluoromount G. Imaging was performed using Elyra 7 microscope (Zeiss). Images were captured using 40x objective. Statistical analysis was performed using Graphpad Prism 9.

#### Immunostaining

Neurons were fixed for 20 min with 4% PFA in PBS at a room temperature. Samples were blocked with 4% donkey serum and 0.2% Tween20 for 1 h. Primary antibodies were diluted in 4% donkey serum and 0.1% Tween20 and applied to samples overnight at 4°C. The following primary antibodies were used chicken MAP2(1:5000, Abcam #ab5392), rabbit GABA (1:1000, SIGMA #A2052), mouse BIN3 (1:500, Invitrogen). Samples were washed with PBS, incubated with fluorescein-conjugated secondary antibodies (Alexa Fluor 647, 546, 488) for 1 h at room temperature, and counterstained with Hoechst for 10 min. Samples were imaged using Elyra 7 microscope (Zeiss).

## Acknowledgements

This research was funded by Aligning Science Across Parkinson’s (ASAP-000301 & ASAP-024434) through the Michael J. Fox Foundation for Parkinson’s Research (MJFF) (to C.R.S with M.B.F., X.D., S.Z., and J.Z.L.), with additional support from the Yale School of Medicine’s Stephen & Denise Adams Center for Parkinson’s Disease Research (C.R.S.), the Yale APDA Center for Parkinson Precision Medicine (C.R.S.), the Lineberger Research Fund (C.R.S.), and the Klarman Cell Observatory (J.Z.L.). C.R.S.’ work was in part supported by NIH R01NS115144 and the U.S. Department of Defense. The collection of brain samples was enabled through NIH P30AG019610 and P30AG072980 (both to T.G.B. and the Arizona Alzheimer’s Disease Center); the Arizona Department of Health Services (T.G.B) and the Arizona Biomedical Research Commission (contracts 4001, 0011, 05-901 and 1001 to the Arizona Parkinson’s Disease Consortium). T.G.B. was in part supported by NIH U24NS072026. M.B.F. was in part supported by NIH R01NS098821 and R35NS132225. We thank Yi Zhong for excellent technical assistance and are grateful to all participants and their families supporting the Banner Brain and Body Donation Program. See additional acknowledgements in the **Supplementary File** for external data sets.

## Author contributions statement

C.R.S. conceived, designed, obtained funding, coordinated, supervised, analyzed, and interpreted the study, and wrote the manuscript with all authors contributing. Z.L. contributed to design and wrote the manuscript, and performed statistical and bioinformatics analyses with contributions from T.W., S.K.S., J.P., X.T., J.Y., X.D. sn-allele-specific expression analysis was conducted by S.K.S., X.A., N.H., and J.Z.L. sn-RNAseq data production was performed by Z.L., I.T., C.B.A., M.S. sn-RNAseq data processing and analysis was performed by J.P., X.T. Stem cell experiments were designed and conducted by S.M. and S-C.Z. *Drosophila* experiments were designed and conducted by V.N. and M.B.F. Brain collection, immunohistochemistry, neuropathological characterization were performed by T.G.B. and G.E.S.. B.W. contributed to manuscript preparation, figures, project organization, data coordination. All authors reviewed, edited, and approved the manuscript prior to submission.

## Data availability

The data, code, protocols, and key lab materials used and generated in this study are listed in a Key Resource Table alongside their persistent identifiers at (Table S11).

## Notes

### Competing Interest Statement

The authors have declared no competing interest.

### Summary of Updates

1. Remove "∽" in the abstract; 2. Change the reuse license to "CC BY".

## References

1. Simuni, T. et al. A biological definition of neuronal alpha-synuclein disease: towards an integrated staging system for research. Lancet Neurol 23, 178–190 (2024).

2. Soto, C. et al. Toward a biological definition of neuronal and glial synucleinopathies. Nat Med 31, 396–408 (2025).

3. Li, M., Ye, X., Huang, Z., Ye, L. & Chen, C. Global burden of Parkinson’s disease from 1990 to 2021: a population-based study. BMJ Open 15, e095610 (2025).

4. Dorsey, E.R., Sherer, T., Okun, M.S. & Bloem, B.R. The Emerging Evidence of the Parkinson Pandemic. J Parkinsons Dis 8, S3–S8 (2018).

5. Mattila, P.M., Rinne, J.O., Helenius, H., Dickson, D.W. & Roytta, M. Alpha-synuclein-immunoreactive cortical Lewy bodies are associated with cognitive impairment in Parkinson’s disease. Acta Neuropathol 100, 285–90 (2000).

6. Hurtig, H.I. et al. Alpha-synuclein cortical Lewy bodies correlate with dementia in Parkinson’s disease. Neurology 54, 1916–21 (2000).

7. Martin, W.R.W. et al. Neocortical Lewy Body Pathology Parallels Parkinson’s Dementia, but Not Always. Ann Neurol 93, 184–195 (2023).

8. Shahmoradian, S.H. et al. Lewy pathology in Parkinson’s disease consists of crowded organelles and lipid membranes. Nat Neurosci 22, 1099–1109 (2019).

9. Xia, Q. et al. Proteomic identification of novel proteins associated with Lewy bodies. Front Biosci 13, 3850–6 (2008).

10. Goh, Y.Y. et al. Multiple system atrophy. Pract Neurol 23, 208–221 (2023).

11. Papp, M.I., Kahn, J.E. & Lantos, P.L. Glial cytoplasmic inclusions in the CNS of patients with multiple system atrophy (striatonigral degeneration, olivopontocerebellar atrophy and Shy-Drager syndrome). J Neurol Sci 94, 79–100 (1989).

12. Poewe, W. et al. Multiple system atrophy. Nat Rev Dis Primers 8, 56 (2022).

13. Irwin, D.J. et al. Neuropathologic substrates of Parkinson disease dementia. Ann Neurol 72, 587–98 (2012).

14. Braak, H. et al. Staging of brain pathology related to sporadic Parkinson’s disease. Neurobiol Aging 24, 197–211 (2003).

15. Beach, T.G. et al. Unified staging system for Lewy body disorders: correlation with nigrostriatal degeneration, cognitive impairment and motor dysfunction. Acta Neuropathol 117, 613–34 (2009).

16. Silbert, L.C. & Kaye, J. Neuroimaging and cognition in Parkinson’s disease dementia. Brain Pathol 20, 646–53 (2010).

17. Williams-Gray, C.H. et al. The distinct cognitive syndromes of Parkinson’s disease: 5 year follow-up of the CamPaIGN cohort. Brain 132, 2958–69 (2009).

18. Schrag, A., Jahanshahi, M. & Quinn, N. What contributes to quality of life in patients with Parkinson’s disease? J Neurol Neurosurg Psychiatry 69, 308–12 (2000).

19. Zhang, W. et al. Aggregated alpha-synuclein activates microglia: a process leading to disease progression in Parkinson’s disease. FASEB J 19, 533–42 (2005).

20. Harms, A.S. et al. MHCII is required for alpha-synuclein-induced activation of microglia, CD4 T cell proliferation, and dopaminergic neurodegeneration. J Neurosci 33, 9592–600 (2013).

21. Liddelow, S.A. et al. Neurotoxic reactive astrocytes are induced by activated microglia. Nature 541, 481–487 (2017).

22. Yun, S.P. et al. Block of A1 astrocyte conversion by microglia is neuroprotective in models of Parkinson’s disease. Nat Med 24, 931–938 (2018).

23. Altay, M.F., Liu, A.K.L., Holton, J.L., Parkkinen, L. & Lashuel, H.A. Prominent astrocytic alpha-synuclein pathology with unique post-translational modification signatures unveiled across Lewy body disorders. Acta Neuropathol Commun 10, 163 (2022).

24. Bryois, J. et al. Genetic identification of cell types underlying brain complex traits yields insights into the etiology of Parkinson’s disease. Nat Genet 52, 482–493 (2020).

25. Nalls, M.A. et al. Identification of novel risk loci, causal insights, and heritable risk for Parkinson’s disease: a meta-analysis of genome-wide association studies. Lancet Neurol 18, 1091–1102 (2019).

26. Kim, J.J. et al. The Parkinson’s Disease DNA Variant Browser. Mov Disord 36, 1250–1258 (2021).

27. Grenn, F.P. et al. The Parkinson’s Disease Genome-Wide Association Study Locus Browser. Mov Disord 35, 2056–2067 (2020).

28. Zeng, B. et al. Single-Nucleus Atlas of Cell-Type Specific Genetic Regulation in the Human Brain. Res Sq (2024).

29. Lopes, K.P. et al. Genetic analysis of the human microglial transcriptome across brain regions, aging and disease pathologies. Nat Genet 54, 4–17 (2022).

30. Humphrey, J. et al. Long-read RNA-seq atlas of novel microglia isoforms elucidates disease-associated genetic regulation of splicing. medRxiv (2023).

31. Haglund, A. et al. Cell state-dependent allelic effects and contextual Mendelian randomization analysis for human brain phenotypes. Nat Genet 57, 358–368 (2025).

32. Fujita, M. et al. Cell subtype-specific effects of genetic variation in the Alzheimer’s disease brain. Nat Genet 56, 605–614 (2024).

33. Bryois, J. et al. Cell-type-specific cis-eQTLs in eight human brain cell types identify novel risk genes for psychiatric and neurological disorders. Nat Neurosci 25, 1104–1112 (2022).

34. Emani, P.S. et al. Single-cell genomics and regulatory networks for 388 human brains. Science 384, eadi5199 (2024).

35. Castonguay, C.-E. et al. A single-cell eQTL atlas of the human cerebellum reveals vulnerability of oligodendrocytes in essential tremor. bioRxiv, 2024.05.22.595233 (2024).

36. Leventhal, M.J. et al. An integrative systems-biology approach defines mechanisms of Alzheimer’s disease neurodegeneration. Nature Communications 16, 4441 (2025).

37. Jang, B. et al. A meta-analysis of single-nucleus expression quantitative trait loci linking genetic risk to brain disorders. Nat Genet 58, 737–747 (2026).

38. Leventhal, M.J. et al. An integrative systems-biology approach defines mechanisms of Alzheimer’s disease neurodegeneration. Nat Commun 16, 4441 (2025).

39. Dong, X. et al. Circular RNAs in the human brain are tailored to neuron identity and neuropsychiatric disease. Nat Commun 14, 5327 (2023).

40. Dong, X. et al. Enhancers active in dopamine neurons are a primary link between genetic variation and neuropsychiatric disease. Nat Neurosci 21, 1482–1492 (2018).

41. Jerber, J. et al. Population-scale single-cell RNA-seq profiling across dopaminergic neuron differentiation. Nat Genet 53, 304–312 (2021).

42. Zhu, Z.H. et al. Integration of summary data from GWAS and eQTL studies predicts complex trait gene targets. Nature Genetics 48, 481-+ (2016).

43. Cannon, J.R. & Greenamyre, J.T. NeuN is not a reliable marker of dopamine neurons in rat substantia nigra. Neurosci Lett 464, 14–7 (2009).

44. Chu, Y. et al. Nurr1 in Parkinson’s disease and related disorders. J Comp Neurol 494, 495–514 (2006).

45. Giambartolomei, C. et al. Bayesian Test for Colocalisation between Pairs of Genetic Association Studies Using Summary Statistics. Plos Genetics 10(2014).

46. Nalls, M. & Blauwendraat, C. Identification of novel risk loci, causal insights, and heritable risk for Parkinson’s disease: a meta-analysis of genome-wide association studies. Lancet Neurology 18, 1091–1102 (2019).

47. Simmons, S.K. et al. Experimental and Computational Methods for Allelic Imbalance Analysis from Single-Nucleus RNA-seq Data. bioRxiv (2024).

48. Edsgard, D., Reinius, B. & Sandberg, R. scphaser: haplotype inference using single-cell RNA-seq data. Bioinformatics 32, 3038–40 (2016).

49. Makarious, M.B. et al. Large-scale rare variant burden testing in Parkinson’s disease. Brain 146, 4622–4632 (2023).

50. Frost, A., Unger, V.M. & De Camilli, P. The BAR Domain Superfamily: Membrane-Molding Macromolecules. Cell 137, 191–196 (2009).

51. Mayle, K.M., Le, A.M. & Kamei, D.T. The intracellular trafficking pathway of transferrin. Biochim Biophys Acta 1820, 264–81 (2012).

52. Betz, W.J., Mao, F. & Smith, C.B. Imaging exocytosis and endocytosis. Curr Opin Neurobiol 6, 365–71 (1996).

53. Ordonez, D.G., Lee, M.K. & Feany, M.B. alpha-synuclein Induces Mitochondrial Dysfunction through Spectrin and the Actin Cytoskeleton. Neuron 97, 108–124 e6 (2018).

54. Collawn, J.F. & Benveniste, E.N. Regulation of MHC class II expression in the central nervous system. Microbes Infect 1, 893–902 (1999).

55. Falcao, A.M. et al. Disease-specific oligodendrocyte lineage cells arise in multiple sclerosis. Nat Med 24, 1837–1844 (2018).

56. Jakel, S. et al. Altered human oligodendrocyte heterogeneity in multiple sclerosis. Nature 566, 543–547 (2019).

57. Kirby, L. et al. Oligodendrocyte precursor cells present antigen and are cytotoxic targets in inflammatory demyelination. Nat Commun 10, 3887 (2019).

58. Grossmann, N. et al. Mechanistic determinants of the directionality and energetics of active export by a heterodimeric ABC transporter. Nat Commun 5, 5419 (2014).

59. Hupp, M.T., Santos, E.N., Heo, D., Call, C.L. & Guttenplan, K.A. Vesicle Fusion in Oligodendrocyte Maturation and Myelination. J Neurosci 44(2024).

60. Baskin, J.M. et al. The leukodystrophy protein FAM126A (hyccin) regulates PtdIns(4)P synthesis at the plasma membrane. Nat Cell Biol 18, 132–8 (2016).

61. Bowser, D.N. & Khakh, B.S. Two forms of single-vesicle astrocyte exocytosis imaged with total internal reflection fluorescence microscopy. Proc Natl Acad Sci U S A 104, 4212–7 (2007).

62. Sole-Domenech, S., Cruz, D.L., Capetillo-Zarate, E. & Maxfield, F.R. The endocytic pathway in microglia during health, aging and Alzheimer’s disease. Ageing Res Rev 32, 89–103 (2016).

63. Kedjouar, B. et al. Molecular characterization of the microsomal tamoxifen binding site. J Biol Chem 279, 34048–61 (2004).

64. Abeliovich, A. & Gitler, A.D. Defects in trafficking bridge Parkinson’s disease pathology and genetics. Nature 539, 207–216 (2016).

65. Kara, E. et al. An integrated genomic approach to dissect the genetic landscape regulating the cell-to-cell transfer of alpha-synuclein. Cell Rep 35, 109189 (2021).

66. Diaz-Ortiz, M.E. et al. GPNMB confers risk for Parkinson’s disease through interaction with alpha-synuclein. Science 377, eabk0637 (2022).

67. Yang, K. et al. White matter changes in Parkinson’s disease. NPJ Parkinsons Dis 9, 150 (2023).

68. Bauernfeind, A.L. & Babbitt, C.C. The predictive nature of transcript expression levels on protein expression in adult human brain. BMC Genomics 18, 322 (2017).

69. Jinn, S. et al. Functionalization of the TMEM175 p.M393T variant as a risk factor for Parkinson disease. Hum Mol Genet 28, 3244–3254 (2019).

70. Zheng, W. et al. pH regulates potassium conductance and drives a constitutive proton current in human TMEM175. Sci Adv 8, eabm1568 (2022).

71. Young, M.D. & Behjati, S. SoupX removes ambient RNA contamination from droplet-based single-cell RNA sequencing data. Gigascience 9(2020).

72. Wu, Y. et al. Pervasive biases in proxy genome-wide association studies based on parental history of Alzheimer’s disease. Nat Genet 56, 2696–2703 (2024).

73. Beach, T.G. et al. Arizona Study of Aging and Neurodegenerative Disorders and Brain and Body Donation Program. Neuropathology 35, 354–89 (2015).

74. Beach, T.G. et al. Unified staging system for Lewy body disorders: correlation with nigrostriatal degeneration, cognitive impairment and motor dysfunction. Acta Neuropathol 117, 613–34 (2009).

75. DelleDonne, A. et al. Incidental Lewy body disease and preclinical Parkinson disease. Arch Neurol 65, 1074–80 (2008).

76. Hughes, A.J., Daniel, S.E., Kilford, L. & Lees, A.J. Accuracy of clinical diagnosis of idiopathic Parkinson’s disease: a clinico-pathological study of 100 cases. J Neurol Neurosurg Psychiatry 55, 181–4 (1992).

77. Hyman, B.T. & Trojanowski, J.Q. Consensus recommendations for the postmortem diagnosis of Alzheimer disease from the National Institute on Aging and the Reagan Institute Working Group on diagnostic criteria for the neuropathological assessment of Alzheimer disease. J Neuropathol Exp Neurol 56, 1095–7 (1997).

78. McKeith, I.G. et al. Diagnosis and management of dementia with Lewy bodies: third report of the DLB Consortium. Neurology 65, 1863–72 (2005).

79. Schroeder, A. et al. The RIN: an RNA integrity number for assigning integrity values to RNA measurements. BMC Mol Biol 7, 3 (2006).

80. Liu, G. et al. Genome-wide survival study identifies a novel synaptic locus and polygenic score for cognitive progression in Parkinson’s disease Nature Genetics in press(2021).

81. Polychronakos, C. & Alriyami, M. Diabetes in the post-GWAS era. Nat Genet 47, 1373–4 (2015).

82. Leek, J.T., Johnson, W.E., Parker, H.S., Jaffe, A.E. & Storey, J.D. The sva package for removing batch effects and other unwanted variation in high-throughput experiments. Bioinformatics 28, 882–3 (2012).

83. Simmons, S.K. et al. Experimental and Computational Methods for Allelic Imbalance Analysis from Single-Nucleus RNA-seq Data. bioRxiv, 2024.08.13.607784 (2025).

84. Di Tommaso, P. et al. Nextflow enables reproducible computational workflows. Nat Biotechnol 35, 316–319 (2017).

85. Kaminow, B., Yunusov, D. & Dobin, A. STARsolo: accurate, fast and versatile mapping/quantification of single-cell and single-nucleus RNA-seq data. bioRxiv, 2021.05.05.442755 (2021).

86. Asiimwe, R. & Alexander, D. STAR+WASP reduces reference bias in the allele-specific mapping of RNA-seq reads. bioRxiv, 2024.01.21.576391 (2024).

87. Li, H. et al. The Sequence Alignment/Map format and SAMtools. Bioinformatics 25, 2078–2079 (2009).

88. Lancelot, M.L.a.R. aod: Analysis of Overdispersed Data. in R package version 1.3, 2012. (Available at http://cran.r-project.org/package=aod, 2023).

89. t Hoen, P.A. et al. Reproducibility of high-throughput mRNA and small RNA sequencing across laboratories. Nat Biotechnol 31, 1015–22 (2013).

90. Price, A.L. et al. Principal components analysis corrects for stratification in genome-wide association studies. Nat Genet 38, 904–9 (2006).

91. Han, B. & Eskin, E. Random-effects model aimed at discovering associations in meta-analysis of genome-wide association studies. Am J Hum Genet 88, 586–98 (2011).

92. Zhou, Y. et al. Metascape provides a biologist-oriented resource for the analysis of systems-level datasets. Nat Commun 10, 1523 (2019).

93. Pinero, J. et al. DisGeNET: a comprehensive platform integrating information on human disease-associated genes and variants. Nucleic Acids Res 45, D833–D839 (2017).

94. Szklarczyk, D. et al. STRING v11: protein-protein association networks with increased coverage, supporting functional discovery in genome-wide experimental datasets. Nucleic Acids Res 47, D607–D613 (2019).

95. Stark, C. et al. BioGRID: a general repository for interaction datasets. Nucleic Acids Res 34, D535–9 (2006).

96. Li, T. et al. A scored human protein–protein interaction network to catalyze genomic interpretation. Nature Methods 14, 61–64 (2017).

97. Shannon, P. et al. Cytoscape: a software environment for integrated models of biomolecular interaction networks. Genome Res 13, 2498–504 (2003).

98. Olsen, A.L. & Feany, M.B. Glial alpha-synuclein promotes neurodegeneration characterized by a distinct transcriptional program in vivo. Glia 67, 1933–1957 (2019).

99. Liu, Y. et al. Directed differentiation of forebrain GABA interneurons from human pluripotent stem cells. Nat Protoc 8, 1670–9 (2013).

