## Supplemental Figures for "Single-cell eQTL mapping reveals convergent glial–neuronal risk architecture in Parkinson’s disease"

**INDEX**

**Supplemental Figure S1……………………………..….……………..………..………3**

**Supplemental Figure S2……………………………..………………..………...…………5**

**Supplemental Acknowledgements…......………………………………...………………......6**

**Supplemental References………..…………………………………...………………......8**

### Supplemental Fig. S1

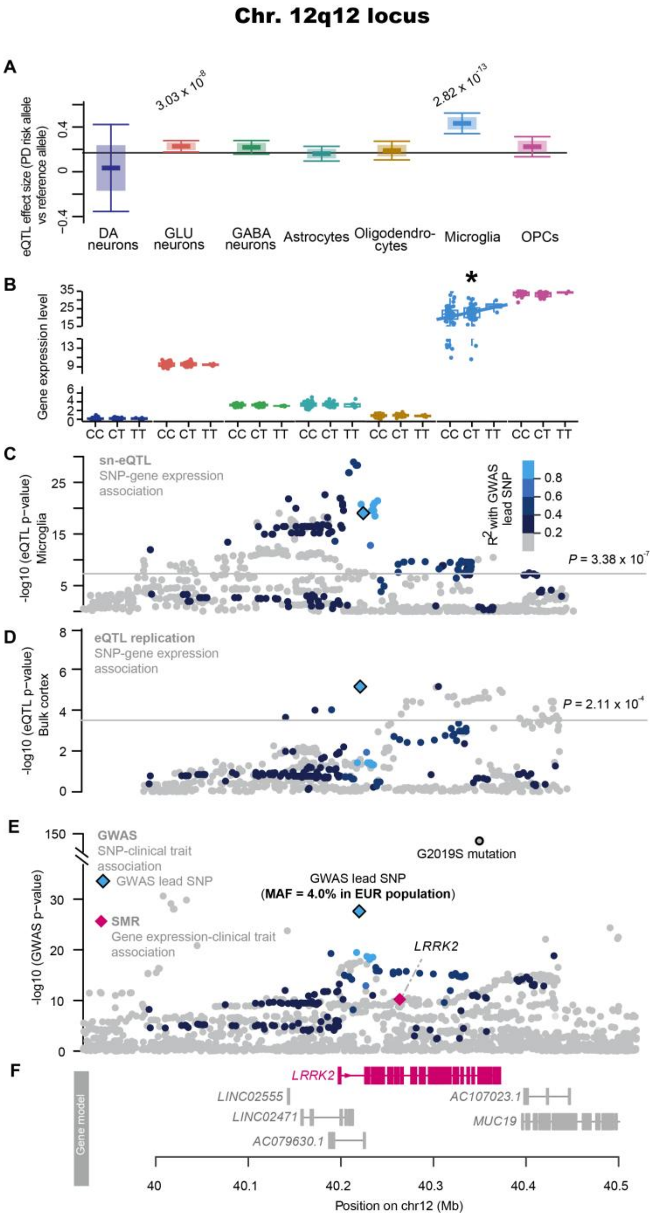


**Fig. S1. *LRRK2* is a quantitative trait gene associated with PD in the Chr. 12q12 GWAS locus that preferentially targets GLU neuron and microglia cells.**

(**A**) Cell type meta-eQTL of *LRRK2* gene. The distribution of meta-eQTL effect sizes and their standard errors across nine brain cell types is represented using box plots. Significant cell types are annotated with their corresponding p-values.

**(B)** Genotype-expression associations in cell types from PD5D and BRAINcode.

**(C-D)** Locus zoom plot shows associations between noncoding variants and *LRRK2* expression for the chr. 12q12 GWAS peak in the GLU neurons **(B)** and microglia **(C)**. Blue color-bar indicates linkage disequilibrium ($R^2$) between GWAS lead variant and other variants (cyan, $R^2$ greater than 0.8). y-axis, negative log10 of $P$ values from cell type meta-eQTL. Bold black outlined diamond, indicates the lead GWAS variant from Ref. [2].

(**E**) Locus zoom plot shows associations between noncoding variants and Parkinson's risk. Gene model showed all of the six genes physically localizing under the GWAS peak. Cell type QTL and SMR analyses prioritized *LRRK2* as functional quantitative trait genes associated with risk of PD (magenta diamonds) out of the six physical candidates localized under this GWAS peak.

**Supplemental Fig. S2**

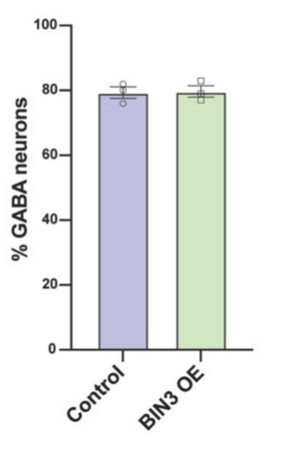


**Fig. S2. *BIN3* overexpression does not affect differentiation into GABAergic neurons.** *BIN3* overexpressing and control hPSCs give rise to similar percentage of GABAergic neurons. Data are presented as mean ± SEM.

### Supplemental Acknowledgements

MSBB: The results published here are in part based on data obtained from the AD Knowledge Portal (**https://adknowledgeportal.org/**). These data were generated from postmortem brain tissue collected through the Mount Sinai VA Medical Center Brain Bank and were provided by Dr. Eric Schadt from Mount Sinai School of Medicine.

HBTRC: The results published here are in whole or in part based on data obtained from the AD Knowledge Portal (https://adknowledgeportal.org). These data were generated from postmortem brain tissue collected through the Harvard Brain Tissue Resource Center and were provided by Dr. Eric Schadt from Mount Sinai School of Medicine.

MayoRNASeq: The results published here are in whole or in part based on data obtained from the AD Knowledge Portal (https://adknowledgeportal.org). The Mayo RNAseq study data was led by Dr. Nilüfer Ertekin-Taner, Mayo Clinic, Jacksonville, FL as part of the multi-PI U01 AG046139 (MPIs Golde, Ertekin-Taner, Younkin, Price). Samples were provided from the following sources: The Mayo Clinic Brain Bank. Data collection was supported through funding by NIA grants P50 AG016574, R01 AG032990, U01 AG046139, R01 AG018023, U01 AG006576, U01 AG006786, R01 AG025711, R01 AG017216, R01 AG003949, NINDS grant R01 NS080820, CurePSP Foundation, and support from Mayo Foundation. Study data includes samples collected through the Sun Health Research Institute Brain and Body Donation Program of Sun City, Arizona. The Brain and Body Donation Program is supported by the National Institute of Neurological Disorders and Stroke (U24 NS072026 National Brain and Tissue Resource for Parkinsons Disease and Related Disorders), the National Institute on Aging (P30 AG19610 Arizona Alzheimers Disease Core Center), the Arizona Department of Health Services (contract 211002, Arizona Alzheimers Research Center), the Arizona Biomedical Research Commission (contracts 4001, 0011, 05-901 and 1001 to the Arizona Parkinson's Disease Consortium) and the Michael J. Fox Foundation for Parkinsons Research.

CommonMind: Data were generated as part of the CommonMind Consortium supported by funding from Takeda Pharmaceuticals Company Limited, F. Hoffmann-La Roche Ltd and NIH grants R01MH085542, R01MH093725, P50MH066392, P50MH080405, R01MH097276, RO1-MH-075916, P50M096891, P50MH084053S1, R37MH057881, AG02219, AG05138, MH06692, R01MH110921, R01MH109677, R01MH109897, U01MH103392, and contract HHSN271201300031C through IRP NIMH. Brain tissue for the study was obtained from the following brain bank collections: the Mount Sinai NIH Brain and Tissue Repository, the University of Pennsylvania Alzheimer's Disease Core Center, the University of Pittsburgh NeuroBioBank and Brain and Tissue Repositories, and the NIMH Human Brain Collection Core. CMC Leadership: Panos Roussos, Joseph Buxbaum, Andrew Chess, Schahram Akbarian, Vahram Haroutunian (Icahn School of Medicine at Mount Sinai), Bernie Devlin, David Lewis (University of Pittsburgh), Raquel Gur, Chang-Gyu Hahn (University of Pennsylvania), Enrico Domenici (University of Trento), Mette A. Peters, Solveig Sieberts (Sage Bionetworks), Thomas Lehner, Stefano Marenco, Barbara K. Lipska (NIMH).

ROSMAP: Study data were provided by the Rush Alzheimer's Disease Center, Rush University Medical Center, Chicago. Data collection was supported through funding by NIA grants P30AG10161 (ROS), R01AG15819 (ROSMAP; genomics and RNAseq), R01AG17917 (MAP), R01AG30146, R01AG36042 (5hC methylation, ATACseq), RC2AG036547 (H3K9Ac), R01AG36836 (RNAseq), R01AG48015 (monocyte RNAseq) RF1AG57473 (single nucleus RNAseq), U01AG32984 (genomic and whole exome sequencing), U01AG46152 (ROSMAP AMP-AD, targeted proteomics), U01AG46161(TMT proteomics), U01AG61356 (whole genome sequencing, targeted proteomics, ROSMAP AMP-AD), the Illinois Department of Public Health

(ROSMAP), and the Translational Genomics Research Institute (genomic). Additional phenotypic data can be requested at www.radc.rush.edu.

GTEx: The Genotype-Tissue Expression (GTEx) Project was supported by the Common Fund of the Office of the Director of the National Institutes of Health, and by NCI, NHGRI, NHLBI, NIDA, NIMH, and NINDS.

UKBEC: UKBEC Acknowledgements

UK Brain Expression Consortium (UKBEC) members and affiliations

Mina Ryten[1,2]

Michael E Weale[1]

John Hardy[2]

Karishma D'Sa[1,2]

Adaikalavan Ramasamy[1,2,3]

Daniah Trabzuni[2,4]

Sebastian Guelfi[2]

Juan A. Botia[2]

Jana Vandrovcova2

Colin Smith[5]

Robert Walker[5]

[1]Department of Medical & Molecular Genetics, King's College London, Guy's Hospital, London, UK

[2]Reta Lila Weston Research Laboratories, Department of Molecular Neuroscience, University College London (UCL) Institute of

Neurology, London, UK

[3]Jenner Institute, University of Oxford, Oxford, UK

[4]Department of Genetics, King Faisal Specialist Hospital and Research Centre, Riyadh, Saudi Arabia

[5]Department of Neuropathology, MRC Sudden Death Brain Bank Project, University of Edinburgh, Edinburgh, UK

### Supplementary references


1. Price, A.L. *et al.* Principal components analysis corrects for stratification in genome-wide association studies. *Nat Genet* 38, 904-9 (2006).
2. Nalls, M.A. *et al.* Identification of novel risk loci, causal insights, and heritable risk for Parkinson's disease: a meta-analysis of genome-wide association studies. *Lancet Neurol* 18, 1091-1102 (2019).
3. Diaz-Ortiz, M.E. *et al.* GPNMB confers risk for Parkinson's disease through interaction with alpha-synuclein. *Science* 377, eabk0637 (2022).
